# Beyond the Visual Word Form Area: Characterizing a hierarchical, distributed and bilateral network for visual word processing

**DOI:** 10.1101/2023.07.11.548613

**Authors:** Raina Vin, Nicholas M. Blauch, David C. Plaut, Marlene Behrmann

**Affiliations:** Neuroscience Institute, Carnegie Mellon University; Department of Psychology, Carnegie Mellon University; Program in Neural Computation, Carnegie Mellon University; Department of Ophthalmology, University of Pittsburgh; Interdepartmental Neuroscience Program, Yale University

**Keywords:** visual word processing, stimulus modulation, VWFA, fMRI, functional connectivity, network analysis, hemispheres, language areas, neural circuit, reading

## Abstract

Although the left hemisphere (LH) Visual Word Form Area (VWFA) is considered the pre-eminent cortical region engaged in visual text processing, other regions in both hemispheres have also been implicated. To examine the entire circuit, using functional MRI data, we defined ten regions of interest (ROI) in each hemisphere that, based on functional connectivity measures, naturally grouped into early vision, high-level vision, and language clusters. We analysed univariate and multivariate responses to words, inverted words, and consonant strings for ROIs and clusters, and demonstrated modulation by text condition bihemispherically, albeit more strongly and in a larger number of regions in the LH. Graph theory analysis revealed that the high-level vision cluster and, specifically, the VWFA was equivalently connected with both early visual and language clusters in both hemispheres, reflecting its role as a mediator in the circuit. Our findings reveal bihemispheric, stimulus-mediated ROI response flexibility but circuit-level connectivity stability, reflecting the complex contribution of a distributed system for word processing.

---

Longstanding data from premorbidly literate patients with acquired reading deficits following brain insult, along with more recent electrophysiological findings from intracranial depth electrodes in humans, have uncovered causal evidence critically implicating the Visual Word Form Area (VWFA) of the left hemisphere (LH) in visual word processing (Behrmann et al., 1998; Cohen et al., 2000; Dehaene and Cohen, 2011; Rosazza et al., 2007; Woolnough et al., 2021, 2022b; A and A, 2018). These findings have been complemented by functional magnetic resonance imaging (fMRI) investigations demonstrating disproportionate selectivity in this region of extrastriate cortex for written words over non-words, faces and objects (Baker et al., 2007; Stevens et al., 2017).

Many studies, however, have pointed out that the VWFA is not a monolithic region, as originally assumed, and that its response properties and connectivity are more complex (Cohen and Dehaene, 2004; Price and Mechelli, 2005; Price and Devlin, 2011). For example, the VWFA is not only responsive to visual stimuli but is also activated by tactile and auditory input, albeit to a lesser extent (Price and Devlin, 2003; Li et al., 2023). Additionally, close scrutiny of the VWFA has revealed subdivisions with a posterior-to-anterior hierarchy of selectivity to individual letters, to bigrams, and to morphemes (Vinckier et al., 2007). Further differentiation has indicated that, within the VWFA, more posterior subregions mediate visual representations, whereas more anterior regions mediate language representations (Caffarra et al., 2021; Yablonski et al., 2019; Grotheer et al., 2021). The activation in these anterior regions predicts reading behavior outside of the scanner (Lerma-Usabiaga et al., 2018) and implements task-specific effects via VWFA-Broca’s connectivity (White et al., 2023). Consistent with the functional distinctions, posterior and anterior regions of the VWFA differ in their micro-architectonic properties (Gomez et al., 2017; Rosenke et al., 2018; Weiner et al., 2017; Lerma-Usabiaga et al., 2018) and in their connectivity profiles. The posterior region, which is involved in feature extraction, is structurally connected to earlier occipital regions and to the intraparietal sulcus via the ventral occipital fasciculus (Yeatman et al., 2014), whereas the anterior region, which integrates information, is connected to the angular gyrus via the posterior arcuate fasciculus and other language regions (Lerma-Usabiaga et al., 2018; Bouhali et al., 2019; Qin et al., 2021).

## The VWFA as a node in a circuit

In addition to uncovering this more complex arrangement of the VWFA, recent investigations have also identified regions beyond the VWFA that are implicated in text processing, including the superior temporal sulcus (STS), the lateral temporal lobe and the posterior inferior parietal lobule (Bouhali et al., 2014; Chen et al., 2019). Also, frontal and parietal regions provide direct top-down modulation to the VWFA (Chen et al., 2019; Qin et al., 2021) that is stronger than that received by any other region of visual cortex (Li et al., 2020).

The inter-relationships between the VWFA and these other regions of interest (ROIs) have been largely established by functional connectivity (FC) analyses. As determined by data from almost 1,000 readers, there are ‘core’ regions such as the insular and frontal opercular cortex, lateral temporal cortex, and early auditory cortex with positive correlations to the VWFA, and, unsurprisingly, responses in these regions are strong predictors of reading performance (Kristanto et al., 2020; for further review see Caffarra et al., 2021). Additionally, FC between the VWFA and Broca’s area, LH Precentral Gyrus (PCG) and Wernicke’s area is observed even during resting state scanning, with the strength of the VWFA-Wernicke’s FC predicting performance on a semantic classification task with words but not with other categories of visual stimuli (Stevens et al., 2017). Furthermore, studies have demonstrated that the pars opercularis, the supramarginal gyrus and the superior temporal gyrus also evince activation in response to orthographic input (Kaestner et al., 2022). Moreover, the strengths of the functional interactions between these different regions can be modulated by the orthographic stimulus, for example, by words versus pseudowords and by the specific task demands with stronger activation during lexical and semantic tasks (Stevens et al., 2017; Planton et al., 2022; White et al., 2023). Of interest too is that FC between VWFA and other regions is heightened for slow compared with fast readers, perhaps indicative of the need of the former to constrain responses further with top-down modulation (Fan et al., 2023). Taken together, these findings suggests that text processing, rather than being solely under the purview of the LH VWFA, is subserved by a widely distributed network of LH regions with a complex system of functional and structural interactions.

In light of the above, the first goal of this paper is to adopt a broad, data-driven lens on the neural correlates of word processing, using fMRI to characterize LH text-selective regions, and to evaluate their connectivity and differential sensitivity to different types of text. In so doing, we also examine whether the multiplicity of activated regions naturally aggregate to form clusters and, using graph theory, uncover the organizational properties of this distributed LH word processing circuit.

## The right hemisphere and word processing

While the predominant focus of most studies has been on the LH VWFA, which is typically co-lateralized with LH dominant language areas (Cai et al., 2010, 2008), recent findings have attested to the engagement of ROIs in the RH as well (Kristanto et al., 2020), not just in individuals with damage to LH VWFA (Liu et al., 2019). Also, the sequence of activation in ventral temporal cortex (VTC) appears similar in both hemispheres, with earlier activation of the posterior fusiform, followed by a spread to medial and lateral VTC, and, then, finally to anterior parts of the brain (Boring et al., 2021). Interestingly, bilaterally, the fusiform gyrus and the STS have greater activation for words over false font stimuli (Woodhead et al., 2011), and for words relative to pseudowords (Bouhali et al., 2019), indicating text type modulation in both hemispheres.

There is, however, a clear hemispheric asymmetry, with greater engagement of the LH than the RH (see Behrmann and Plaut (2020) for review), and earlier latency for LH than RH word-selective responses (Boring et al., 2021). One recent proposal suggests that an initially bilateral circuit encompassing posterior left and right VWFA-1 subregions (see White et al. (2019) for these divisions) converge to send hemifield-specific input to left VWFA-2, which displays sensitivity to lexical properties of only one attended word, consistent with the observed behavioral processing bottleneck (White et al., 2019). Also, close inspection reveals that the hemispheric profiles diverge as a function of format familiarity: horizontally and vertically presented words yield the same activity in the LH VWFA, whereas the RH is more activated by the latter (Cai et al., 2010), perhaps as a result of RH mental rotation computations (Gauthier et al., 2002). The second goal of the current study, then, is to expand our investigation beyond the LH and undertake a similar, comprehensive exploration of the RH, with a focus on the regions engaged in word perception, their modifiability by different text conditions, and their relationships to the LH homologous regions.

## The current study

It is clear that the neural mechanisms of text processing extend beyond the LH VWFA region with additional regions being identified across a host of different studies. Here, we characterize the organization of the entire circuit that is functionally implicated in text processing. Beyond demarcating many ROIs and the pairwise relationship between them, we evaluate the hierarchical organization that emerges from clustering regions with similar activation profiles. We measure the univariate selectivity, multivariate response (decoding) and functional connectivity at both the ROI and cluster level. Last, we measure the node strength of each ROI and cluster, to provide a broad-scale characterization of text-selective cortex.

The examination of a widespread network fits with neuroscience proposals which call for scientific inquiry to account for integration of widespread network processes, and their dynamics and flexibility (Bassett and Sporns, 2017). In particular, we hone in on the following multi-part questions: (1) What are the primary regions that comprise the visual word processing network? (2) What is the nature of the connectivity profile among these ROIs? Are there clusters of ROIs based on their particular functional properties? Are there differences in the activation profile at the ROI and cluster level as a function of stimulus type? (3) What are the circuit-wide properties of the visual word processing network within- and between-hemispheres as measured by graph theoretic analyses?

## Methods

### Participants

A sample of 28 right-handed native English speakers (mean age: 22.1 yrs, 20 female), with normal or corrected-to-normal visual acuity, participated in this study. Participants were right-handed, native English speakers (not multilingual in childhood), with no significant neurological or psychiatric history, claustrophobia, history of metal in body, or any other counter-indication for MRI. The group had a mean score on the Edinburgh Handedness Inventory (Oldfield, 1971) of 83.7, consistent with right-handedness. Participants completed a 1-hour imaging session consisting of functional, structural, and diffusion scans, and were paid for their participation. The diffusion data form part of a different study and are not discussed in this paper.

The procedures used in this study were approved by the Institutional Review Board of Carnegie Mellon University. All participants gave informed consent prior to their scanning session.

### fMRI experiment

All functional images were acquired on a Siemens Prisma 3T scanner with a 64-channel head coil at the Carnegie Mellon and University of Pittsburgh BRIDGE Center. Five runs of functional imaging data (TE=30 ms, TR=2000 ms) were collected, at isotropic resolution of (2mm)3. Each run lasted approximately 5 minutes and contained several mini-blocks of multiple images from a given category presented rapidly at fixation while the subjects performed a 1-back stimulus identity task. We used a modified version of the *fLoc* category localizer (Stigliani et al., 2015), with 6 categories: words, inverted words, non-word letter strings, faces, inverted faces and objects. Faces were ambient adult face images with natural variability in expression and viewpoint. Objects were a mixture of cars and guitars presented at a variety of orientations. Words were common 5-letter words (see Appendix E for full list), presented in a variety of fonts. Non-words were consonant strings derived from the original words with replacement of vowels with random consonants, but were otherwise identical to the word stimuli in font and presentation. Inverted words and faces were the stimuli from the words and faces categories, with images inverted by a 180 deg rotation. Faces and objects were taken from the original stimuli from Stigliani et al. (2015). Luminance histograms of all stimuli were matched using the histMatch function of the SHINE toolbox (Willenbockel et al., 2010). All stimuli appeared on scrambled backgrounds, and there were 104 per category. Rest mini-blocks were interleaved with the category mini-blocks, with the same frequency and duration as stimulus mini-blocks. Each mini-block lasted 6s, and contained 12 stimuli from the category of interest (or 0 in the control/no-stimulus condition), each presented for 400ms with a 100ms inter-stimulus interval (ISI). Subjects pressed a button on an fMRI compatible glove when detecting a repeat of the previous image (1-back stimulus identity task) which appeared randomly once per run. Participants had 1s to respond from the offset of the repeated image. The order of mini-blocks was fully counterbalanced across categories, such that a mini-block of each category preceded a mini-block of each other category exactly once in a run, resulting in 6 mini-blocks per category, and a run length of 306s. At the end of each run, participants received feedback about their performance.

### Functional data pre-processing

The functional data were preprocessed using *fMRIPrep* 1.4.1 (Esteban et al., 2018b,a), which is based on *Nipype* 1.2.0 (Gorgolewski et al., 2011, 2018). Many internal operations of *fMRIPrep* use *Nilearn* 0.5.2 (Abraham et al., 2014), mostly within the functional processing workflow. For more details of the pipeline, see the section corresponding to workflows in *fMRIPrep*’s documentation. Here, we quote the methods output directly from *fMRIPrep* as applied in our experiment, a procedure encouraged by the *fMRIPrep* authors to ensure accuracy of the methods description. Slight modifications were made to remove the details regarding estimation of unused confounds.

For each of the 5 BOLD runs per participant, the following preprocessing was performed. First, a reference volume and its skull-stripped version were generated using a custom methodology of *fMRIPrep*. A deformation field to correct for susceptibility distortions was estimated based on two echo-planar imaging (EPI) references with opposing phase-encoding directions, using 3dQwarp (Cox and Hyde, 1997) (AFNI 20160207). Based on the estimated susceptibility distortion, an unwarped BOLD reference was calculated for a more accurate co-registration with the anatomical reference. The BOLD reference was then co-registered to the T1w reference using bbregister (FreeSurfer) which implements boundary-based registration (Greve and Fischl, 2009). Co-registration was configured with nine degrees of freedom to account for distortions remaining in the BOLD reference. Head-motion parameters with respect to the BOLD reference (transformation matrices, and six corresponding rotation and translation parameters) were estimated before any spatiotemporal filtering using mcflirt (FSL 5.0.9, Jenkinson et al., 2002). The BOLD time-series were resampled onto their original, native space by applying a single, composite transform to correct for head-motion and susceptibility distortions. These resampled BOLD time-series will be referred to as *preprocessed BOLD in original space*, or just *preprocessed BOLD*. Several confounding time-series were calculated based on the *preprocessed BOLD*: framewise displacement (FD), DVARS and three region-wise global signals. FD and DVARS were calculated for each functional run, both using their implementations in *Nipype* (following the definitions by Power et al., 2014). Additionally, a set of physiological regressors was extracted to allow for component-based noise correction (*CompCor*, Behzadi et al., 2007). Principal components were estimated after high-pass filtering the *preprocessed BOLD* time-series (using a discrete cosine filter with 128s cut-off) for the anatomical *CompCor* variants (aCompCor).

This subcortical mask was obtained by heavily eroding the brain mask, which ensures that it does not include cortical GM regions. For aCompCor, components were calculated within the intersection of the aforementioned mask and the union of CSF and WM masks calculated in T1w space, after their projection to the native space of each functional run (using the inverse BOLD-to-T1w transformation). Components were also calculated separately within the WM and CSF masks. The head-motion estimates calculated in the correction step were also placed within the corresponding confounds file. The confound time series derived from head motion estimates and global signals were expanded with the inclusion of temporal derivatives and quadratic terms for each (Satterthwaite et al., 2013). All resamplings can be performed with *a single interpolation step* by composing all the pertinent transformations (i.e. head-motion transform matrices, susceptibility distortion correction when available, and co-registrations to anatomical and output spaces). Gridded (volumetric) resamplings were performed using antsApplyTransforms (ANTs), configured with Lanczos interpolation to minimize the smoothing effects of other kernels (Lanczos, 1964). Non-gridded (surface) resamplings were performed using mri_vol2surf (FreeSurfer).

### fMRI General Linear Model (GLM)

fMRI data were analyzed using SPM12. For each subject, a general linear model was specified with an event-related design. Specifically, each mini-block was modeled with its onset and a duration of 6 seconds. The design matrix thus contained 6 stimulus regressors, one per condition. Stimulus timecourses were convolved with a canonical hemodynamic response function. A reduced set of FMRIPREP-generated confounds was retained for nuisance regression. Specifically, we retained 6 motion parameters (X, Y, Z motion and rotation), the top 6 principal components of the aCompCor decomposition, and the framewise-displacement, yielding a total of 13 nuisance regressors, along with a runwise mean regressor. Finally, an autoregressive-1 model was used within SPM12 to reduce the effects of serial correlations. We fit voxels only within the brain mask defined by *fMRIPrep*. Estimation of the GLM resulted in a beta-weight for each stimulus condition and run, which were used for subsequent univariate and multivariate analyses.

### Functional selectivity

SPM12 was used to compute whole-brain statistical maps indexing the selectivity for each condition of interest compared to the others. These contrasts were balanced in total weight across numerator and denominator. We computed selectivity for words, letter strings, and inverted words as contrasts against other non-text conditions, e.g. (words > (.25*(faces + inverted faces) + .5*objects)). As words, letter strings, and inverted words evinced similar large-scale responses in ventral regions, we also analyzed a text selectivity contrast: (words + letter strings + inverted words)/3 > (.25*(faces + inverted faces) + .5*(objects)). For some analyses, statistical maps used only the even or odd runs of the experiment so as to assess the reliability of the statistical maps.

### Region of Interest (ROI) definition

We used the *Glasser* atlas to divide each hemisphere into 180 anatomical parcels (Glasser et al., 2016) from which we identified ROIs mentioned in existing papers including the VWFA, Inferior Frontal Gyrus (IFG) and the Peri-Sylvian Language (PSL) region. We also included the IFG orbitofrontal region (IFGorb), the Inferior Parietal Sulcus (IPS), and visual areas V1, V2, and V3, as well as the Superior Temporal Sulcus and Gyrus (abbreviated as STS+G) and the Precentral Gyrus (PCG), as referenced by Stevens et al. (2017). A surface-based alignment was performed using the *Glasser* atlas aligned to the *fsaverage* subject space; after alignment to each subject’s surface, the surface-aligned subject-specific atlas was projected into the cortical ribbon in order to acquire a volumetric atlas consisting of cortical voxels, allowing for analyses in native volumetric space. Based on previous literature (supplementary information, Glasser et al. (2016)), we defined each of these anatomical regions of interest in each hemisphere as being composed of one or more Glasser parcels, as listed in Table 1 and illustrated in Figure 2. For our multivariate analyses, we used a purely anatomical definition of all regions, as defined in Table 1; as the VWFA is typically defined functionally, for clarity we refer to the anatomically defined region as ‘VWFA_a’. For the univariate selectivity, functional connectivity and node strength analyses, we functionally localized each of the ten ROIs in each hemisphere, including the VWFA, within the anatomical bound of their corresponding Glasser parcels from Table 1 (discussed in detail in the following sections). Localization was performed using the statistical map with the text selectivity contrast, as described above, and a localization threshold of *p* < 0.1 (we used a lax threshold so as to capture and measure signal even from ROIs in the RH, some of which had much weaker selectivity than their homologous counterparts in the LH).

**Table 1:**
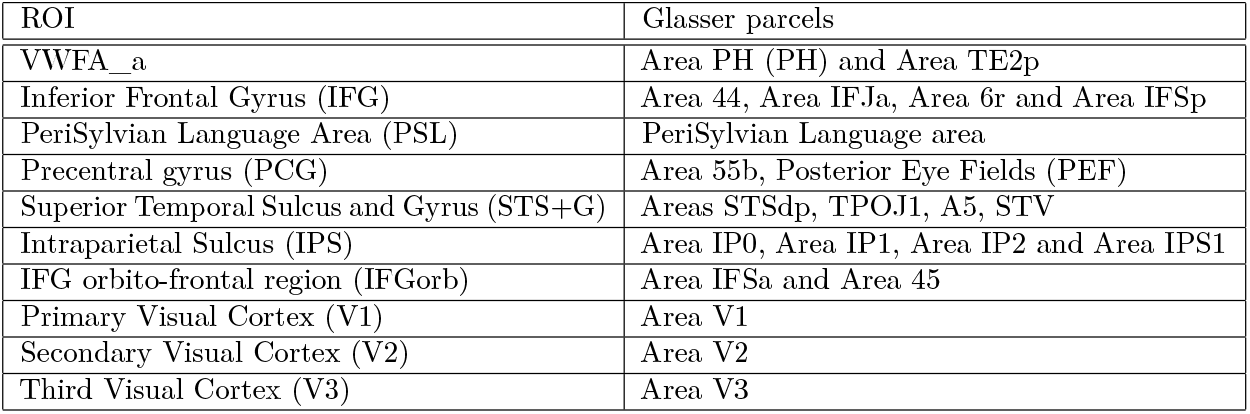
Anatomical search spaces for defining each ROI. Each ROI is composed of one or more anatomical parcels from the Glasser atlas (Glasser et al., 2016). Areas V1-V3 were not aggregated but still constitute ROIs.

**Figure 1:**
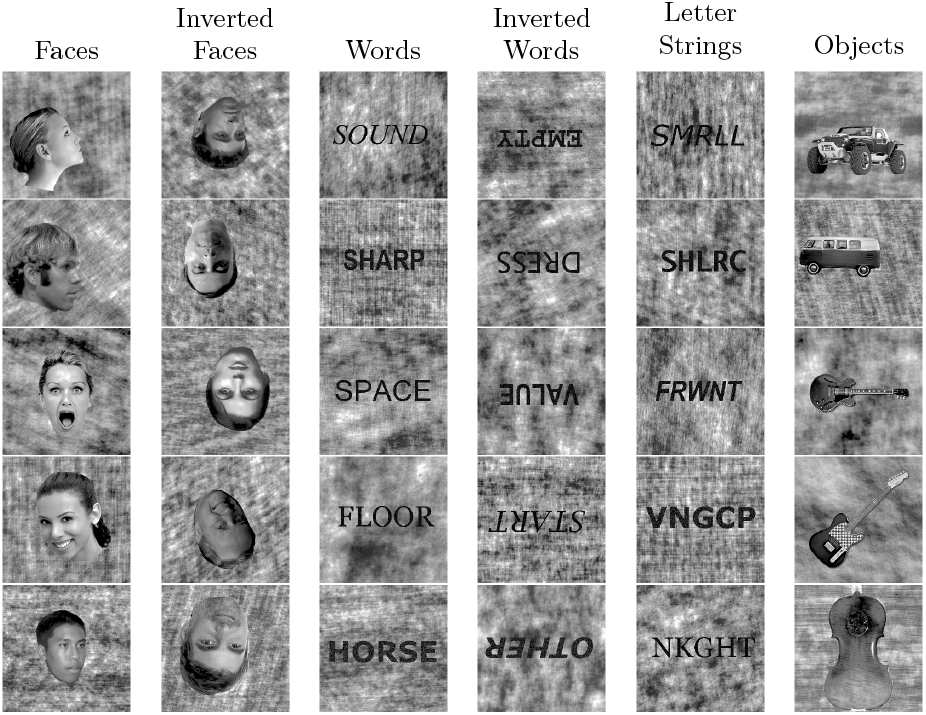
Examples of each stimulus category

**Figure 2:**
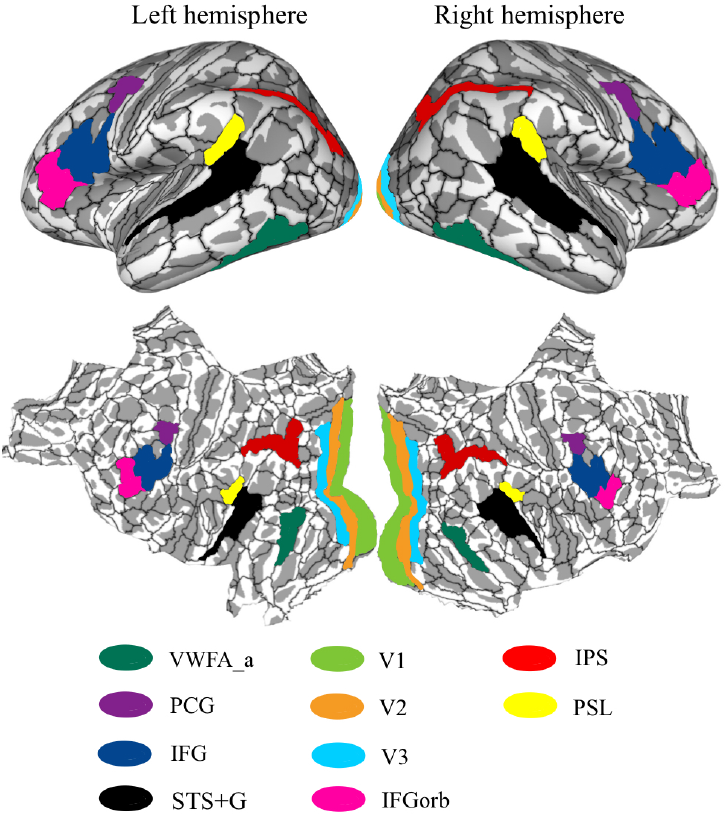
Defining the ROIs involved in visual text processing. ROI-level parcellation using sets of Glasser parcels, viewed on lateral and flatmap views of the cortical surface of both hemispheres.

### Network cluster definition

At the outset, we examined whether, among these multiple ROIs, there was any inherent organization in which subsets of ROIs naturally clustered based on similarity of response profile. To assess this, we took the mean pairwise FC matrix consisting of the correlation values of the time course between ROI pairs, plotted in Figure 3B, and subjected this to the Paris agglomerative hierarchical clustering algorithm (Bonald et al., 2018). Note that this clustering was performed only for the Words condition, which is typically used to identify the VWFA. The dendrogram of the clustering analysis is plotted in Figure 3A (as shown in Appendix C, the same three clusters emerge when the clustering analysis is performed on the other text stimuli). The “Early vision” ROIs (V1, V2, V3) split from all other ROIs, which then split into two clusters, referred to here as the “High-level vision” and “Language” clusters. The Early vision cluster splits into V1 and V2/V3, followed by the latter cluster splitting into separate left and right hemispheric clusters. The High-level vision cluster splits into sets of homologous ROIs (e.g. left and right VWFA and, left and right IPS) and, last, the Language cluster (STS+G, PSL, IFG, IFGorb, PCG) is strongly divided into separate clusters of LH and RH ROIs, with one exception, which is that RH STS+G fell in the otherwise LH-dominant language cluster. To ensure that this clustering was not dependent on the particular agglomerative approach, we repeated this procedure using the Louvain algorithm, with default resolution parameter (*γ* = 1). The output confirmed our initial Paris clustering. We take as our structure the clustering at the *highest* level (see blue lines), with three clusters–Early vision, High-level vision and Language–probed separately in the left and right hemispheres. Given this organizational substructure, below we report the statistical analyses at the single ROI level, pairwise ROI level using FC (both within and between hemispheres) and at the level of the three-cluster parcellation in each hemisphere (by resampling the data using the parcel of the entire cluster rather than simply averaging the activation of its sub-ROIs; see details below in each section).

**Figure 3:**
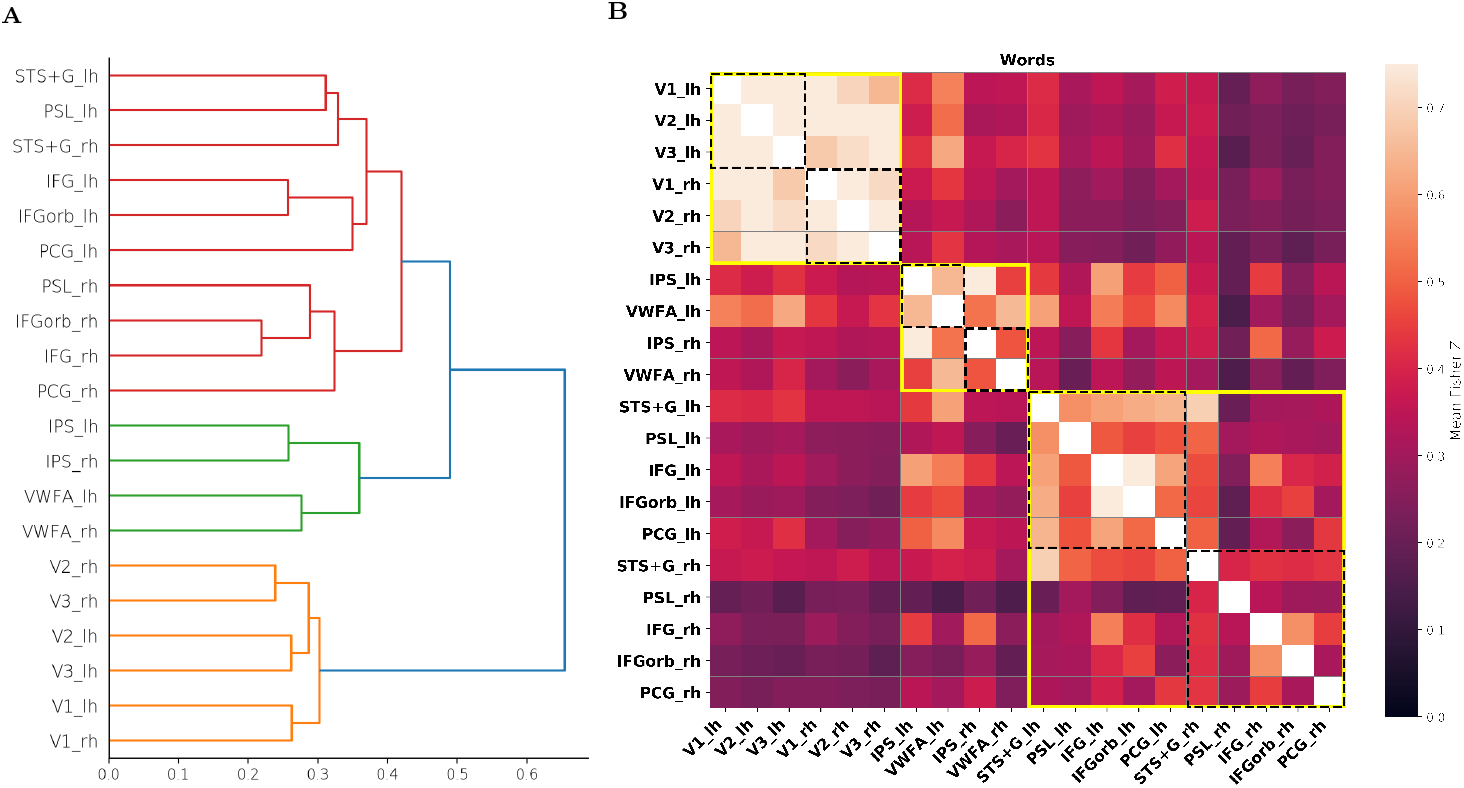
Constructing clusters from patterns of functional connectivity. **A**. Dendrogram showing agglomerative hierarchical clustering result using the Paris algorithm. The three colored clusters correspond to the three clusters analyzed throughout the paper (orange: Early vision, green: High-level vision, red: Language). **B**. Pairwise functional connectivity matrix. Each cell plots the mean Fisher Z score across subjects for functional connectivity (correlation of mean parcel time-series) between the corresponding ROI pair during viewing of Words (see Methods for further details). Yellow boxes indicate the clusters revealed through the clustering algorithm (see **A**., and text for further details). Dashed black boxes show each hemisphere within each of the clusters.

### ROI and cluster-level selectivity

We identified the voxels that were selective for text, and measured their selectivity using a localization approach with both halfs of the data i.e., first using the even runs to functionally localize the voxels and the odd runs to measure selectivity, and second using the odd runs to localize the voxels and the even runs to measure selectivity. For individual ROIs, the anatomical search spaces and contrasts were defined as in Table 1. For cluster-level selectivity, the search space consisted of all of the voxels within all of the ROIs that constitute the given cluster, based on the results of the clustering algorithm (Figure 3A). For each ROI and cluster, we computed the mean of the selectivity using the even runs and the odd runs, which we refer to as *localized selectivity*.

To capture both the strength and size of the selective region within each anatomical parcel, we used the metric of *summed selectivity* following localization of candidate voxels. Using either the even or odd runs of fMRI data, for each ROI, a mask was generated within the respective anatomical search space corresponding to voxels meeting the selectivity criterion (*p* < 0.01 unc.). Then, using the other subset of runs to compute selectivity within these voxels, the t-statistics were summed across voxels within this mask. We computed summed selectivity for each ROI, hemisphere, and stimulus condition.

### Multi-Voxel Pattern Analysis (MVPA)

For the MVPA decoding analyses, we used the beta weights from the GLM that modeled each stimulus condition (i.e., words, inverted words, letter strings, objects, faces and inverted faces) over each of 5 runs. To account for voxel-level uncertainty in beta-estimates, each beta weight was compared to 0 using a one-hot *t*-test, and the *t*-statistics were retained as the voxel-level features, one per stimulus condition per run. A cross-validated decoding was then performed between pairs of text conditions (words, inverted words and letter strings). Additionally, we performed decoding of each text condition against objects. Decoding was done at the ROI level by using the complete set of voxels in each ROI, and at the cluster level by using all voxels in the cluster. A linear classifier with a Ridge (i.e. L2) penalty was used, with the optimal regularization strength determined via internal cross-validation over the training runs. Each run served as the test run for a classifier trained on the other four runs, and the mean classification accuracy over runs was retained.

### Stimulus-based functional connectivity

Stimulus-based FC analyses were conducted on the cortical surface. For each participant, we used 3 out of the 5 runs to functionally localize the selective voxels within the anatomically defined ROIs from Table 1. The activation time series of all voxels within the constituent Glasser parcels within each of the ROIs were then averaged for each of the remaining 2 runs. The time series of the 2 runs were then concatenated into a single time series. To study the visual word processing network, we extracted the time series that corresponded to the peak of the hemodynamic response function (the time-frame of 6*s* to 10*s* after condition onset was used to extract the peak, with TR = 2*s*) elicited by each stimulus condition (i.e., the 1-back perception tasks on Words, Inverted Words and Letter Strings). Using the same confounds used in our GLM analysis, we extracted the residual denoised BOLD time series of each voxel for each stimulus condition. Finally, the mean denoised time series of pairs of ROIs were correlated using Pearson’s *r*, for each stimulus condition. All correlations were further transformed using Fisher’s *z* before being analyzed. At the cluster-level, the functional connectivity between all pairs of constituent ROIs was averaged for each of the three stimulus types, in each hemisphere.

### Graph theory

Each of the ROIs was considered a ‘node’ in the visual word processing network. We used the graph-theoretic metric of node strength to analyze the circuit-level properties of the ROIs. Node strength was computed as the sum of the weights of the edges connected to a given ROI; this was performed either globally (i.e. across all edges), or within a cluster (only including ipsilateral edges and the edge to the homologous contralateral ROI). The weights of the edges between each pair of ROIs was measured as the degree of absolute FC between them.

### Statistical tests

In each analysis, for each dependent measure (e.g., univariate selectivity or multivariate decoding accuracy), repeated-measures ANOVAs were conducted with all relevant within-subject factors. Thereafter, to decompose any observed interactions, we used conservative post-hoc Tukey Honest Significant Difference (HSD) correction on the highest order interaction. The HSD measure is a single-step multiple comparison procedure that permits identifying those means that are significantly different from each other. We restricted the number of pairwise comparisons in the HSD calculation to the differences of interest and used *p* < 0.01 to be highly conservative. Any two means that differed from each other by a value greater than the computed critical Tukey HSD value were considered statistically significant. In some analyses, we adopted additional procedures and these are described in the relevant sections.

## Results

### Selectivity as a function of ROI, cluster, hemisphere and stimulus type

First, we assessed the selectivity for textual stimuli (words, inverted words and letter strings) in each of the anatomical ROIs in each hemisphere. To do so, we computed the localized selectivity of each of the ROIs for each stimulus type against the other non-text conditions (see Methods for details). The significance was determined using one-sample *t*-tests against the null distribution across subjects (Figure 4A).

**Figure 4:**
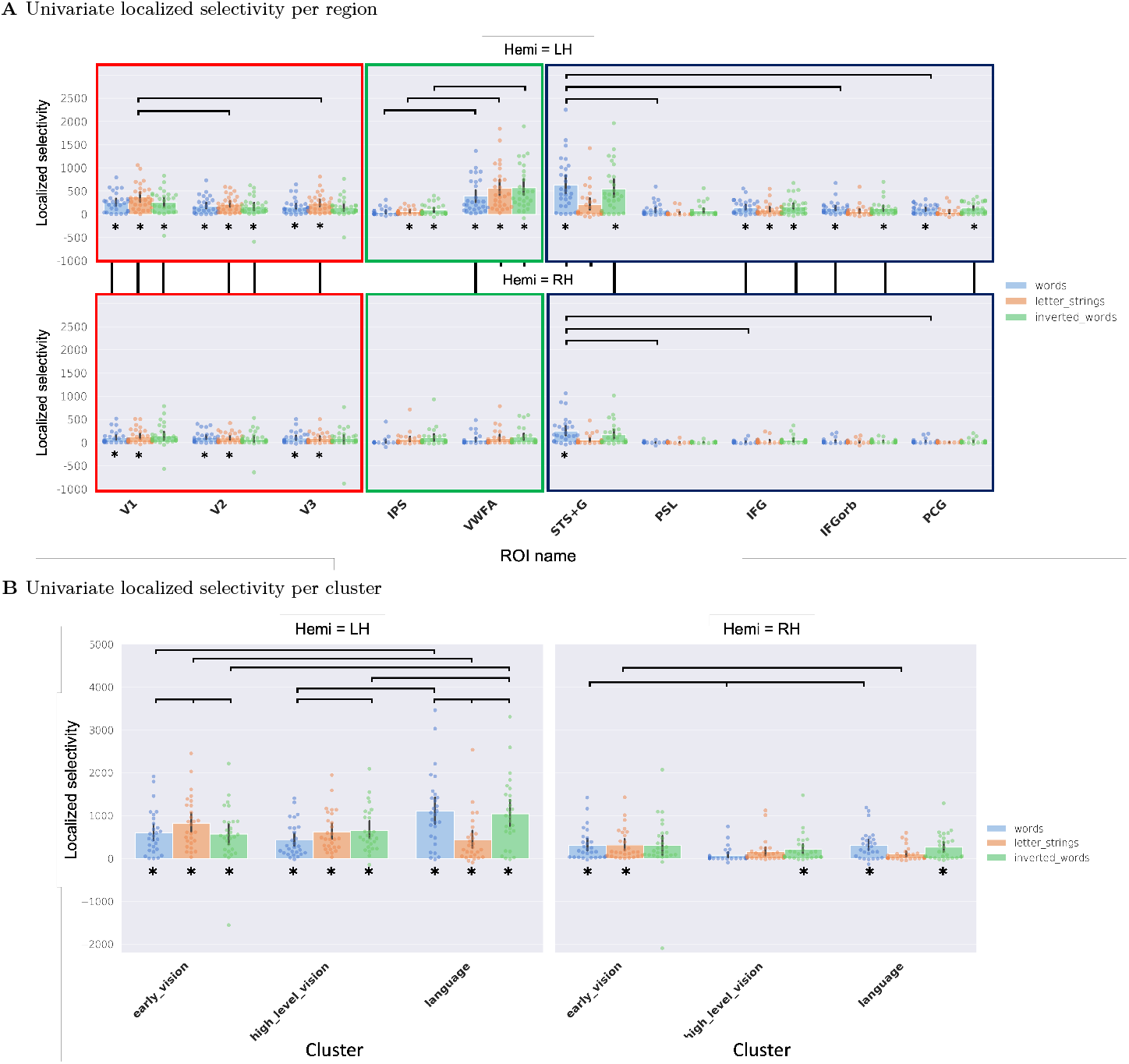
Effects of stimulus type and hemisphere on univariate localized selectivity across ROIs and clusters. **A**. Localized selectivity of each ROI for each of the three stimulus types in each hemisphere (for convenience, we group the ROIs according to their cluster membership so this can be appreciated as well). Horizontal black lines indicate the top 30% of all significant pairwise differences across ROIs and stimulus types in each hemisphere. Vertical black lines indicate all significant between-hemisphere pairwise differences across homologous ROI-stimulus combinations that survived correction. **B**. Localized selectivity per cluster for each stimulus condition. Horizontal black lines indicate all significant within-hemisphere pairwise differences (across clusters and stimulus types). All homologous (i.e., for the same cluster-stimulus combination) between-hemisphere comparisons are significant (see text). In both **A** and **B**, asterisks indicate ROI/cluster×stimulus combinations in the LH and RH that have significantly greater selectivity than the null (i.e., 0) (computed using one-sample *t*-tests and corrected for multiple comparisons). Dots show scores for individual subjects, bars show the average across subjects, and black error lines indicate the 95% confidence interval of scores across subjects. Significance of all pairwise differences was computed with Tukey’s Honest Significant Difference (HSD) test and *p* < 0.01.

We then conducted a repeated measures ANOVA with the localized selectivity as the dependent measure, and ROI, hemisphere and stimulus type as within-subject factors. As evident in Figure 4A, there was a significant three-way interaction of ROI×Hemisphere×Stimulus-type (F(18, 486) = 11.32, *p* < 0.001, *η*^2^ = 0.30). The decomposition of the three-way interaction revealed many pairwise differences both within and between hemispheres. The set of all pairs of ROIs that differed significantly from each other is listed in Appendix A. In Figure 4A, we include horizontal black lines only between the top 30% (a large enough number to provide a representative sample of the results while making the figure details still visible) of all significant pairwise differences across ROIs and stimulus types within each hemisphere, as well as all significant comparisons between hemispheres. As shown by the vertical black lines joining the two hemispheres in the figure, in general, there was greater selectivity for at least one text-type in the LH than in the RH for all ROIs including V1-V3, with the exception of IPS and PSL. The PSL was the only ROI in the LH that did not exhibit significant selectivity to any of the three stimulus types. Across the other ROIs in the Language cluster, the LH selectivity advantage held to a greater extent for words followed by inverted words and then by letter strings. In contrast, the Early vision and High-level vision ROIs exhibited a stronger LH selectivity advantage for letter strings and inverted words. In the LH, the VWFA (particularly for inverted words and letter strings) and STS+G (particularly for words and inverted words) exhibited the greatest selectivity (greater than that of the early visual ROIs – V1, V2 and V3). In the RH, in addition to V1, V2 and V3 (which were similarly selective to words, inverted words and letter strings), the STS+G was also significantly activated (particularly in response to words), with the highest average selectivity across the three stimulus types compared with all other ROIs in the RH.

As can be inferred from the three-way interaction, there were significant two-way interactions of ROI × Hemisphere (F(9, 243) = 13.14, *p* < 0.001, *η*^2^ = 0.33), reflecting greater localized selectivity for many, albeit not all, ROIs in the LH compared with RH; and ROI ×Stimulus (F(18, 486) = 16.75, *p* < 0.001, *η*^2^ = 0.38), indicating greater localized selectivity for words and inverted words compared to letter strings in STS+G in both the LH and RH. The Hemisphere×Stimulus interaction was not significant (F(2, 54) = 1.08, *p* = 0.35, *η*^2^ = 0.04)). The main effects of ROI (F(9,243) = 13.96, *p* < 0.001, *η*^2^ = 0.34) and Hemisphere (F(1, 27) = 78.46, *p* < 0.001, *η*^2^ = 0.74) reached statistical significance, while that of Stimulus (F(2, 54) = 2.91, *p* = 0.06, *η*^2^ = 0.1) trended towards significance.

We also sought to study selectivity at the level of the cluster. This provides a higher-level view of text activation. As with the ROI analysis above, we began by computing the functionally localized selectivity of each of the clusters in each hemisphere for all three textual stimulus types (words, inverted words and letter strings) against the non-text conditions, followed by one-sample *t*-tests against the null distribution across subjects (Figure 4B). In the LH, all three clusters (Early vision, High-level vision and Language) exhibited significant selectivity for all three stimulus types. In the RH, the Early vision cluster was significantly selective for words and letter strings, the High-level vision cluster was selective for inverted words, and the Language cluster was selective for words and inverted words. The significant selectivity for the clusters, for example, in High Level vision, but not in the individual ROIs making up this cluster suggests that there is activation that is cumulative but is not discernible at the level of the ROI.

To measure differences in selectivity between the clusters and the corresponding effect of stimulus type and hemisphere, we repeated the same ANOVA as above but with selectivity at the level of the cluster as the dependent variable. As shown in Figure 4B, there was a significant three-way interaction (F(4, 108) = 20.68, *p* < 0.001, *η*^2^ = 0.43). Post-hoc testing revealed greater LH than RH cluster activation for all three stimulus types for all three clusters (no lines on figure as all are significant). The black horizontal lines in Figure 4B display all within-hemisphere pairwise comparisons whose difference surpassed the HSD threshold (the full set of all pairs that differed significantly from each other is listed in Appendix A). In the LH, all three clusters exhibited significant modulation of their localized selectivity by stimulus type. In particular, the Early vision cluster displayed the highest selectivity for letter strings, the High-level vision cluster had similar selectivity for inverted words and letter strings (both of which were greater than the selectivity for words), and the Language cluster had substantially stronger selectivity for words and inverted words compared to letter strings. Modulation of cluster-level selectivity by stimulus type was less prominent in the Early vision and Language clusters in the RH, even though all three clusters were significantly selective for at least one text condition. There were also several cross-cluster differences as a function of stimulus type in both the LH and RH.

Along with the three-way interaction, there was a significant two-way interaction of Cluster×Hemisphere (F(2, 54) = 3.62, *p* = 0.03, *η*^2^ = 0.12), reflecting the higher localized selectivity for clusters in the LH than RH, especially for the High-level vision and Language clusters. There was also an interaction of Cluster×Stimulus (F(4,108) = 20.80, *p* < 0.001, *η*^2^ = 0.44), with the High-level vision cluster exhibiting greater selectivity for Inverted Words and Letter Strings compared to Words in both hemispheres. The Language cluster, on the other hand, was bihemispherically more selective for Words and Inverted Words compared to Letter Strings (likely driven primarily by the selectivity of the STS+G). Surprisingly, the interaction of Hemisphere × Stimulus was not significant (F(2,54) = 1.08, *p* = 0.35, *η*^2^ = 0.04), reflecting equivalent bihemispheric sensitivity to stimulus type. Last, there was a main effect of Hemisphere (LH > RH: F(1, 27) = 78.46, *p* < 0.001, *η*^2^ = 0.74), and an effect of Stimulus (F(2, 54) = 2.91, *p* = 0.06, *η*^2^ = 0.09).

In summary, the univariate selectivity analysis at both the ROI and cluster level brings out clear differences between the hemispheres. Generally higher localized selectivity was revealed as well as increased stimulus-based modulation in the LH over the RH (albeit with Words and Letter Strings differing to a greater extent than Words and Inverted Words in most Language ROIs). At the ROI level, the LH VWFA and STS+G had the highest average selectivity among all other ROIs in the LH, while the PSL had the lowest average selectivity. In the RH, the STS+G was the most selective ROI, particularly for the Words stimulus, and was closely followed by V1, V2 and V3 (which were relatively similarly selective to all three stimulus types). Notably, while V1-V3 demonstrated text selectivity in the RH, LH selectivity was typically significantly stronger, demonstrating that the LH preference for texts extends even into early visual areas. Selectivity at the cluster level was also significantly higher in the LH than RH, and was modulated by stimulus type. Particularly in the LH, there was greater selectivity in the High-level vision cluster for inverted words compared to words (intermediate selectivity for letter strings). The Language cluster, on the other hand, had similar selectivity for words and inverted words, both of which elicited significantly higher selectivity than letter strings. Despite this hemispheric advantage, the grouping of ROIs into clusters revealed significant selectivity for at least one text condition in each of the 3 RH clusters. For language, this appears to be driven predominantly by the RH STS+G, which demonstrated significant word selectivity, and trending significance for inverted words. For high-level vision in the RH, the trending selectivity for inverted words in IPS and VWFA became significant at the level of the cluster, implying a weak but significant involvement of the RH within high-level selective visual processing of text.

### Multivariate information as a function of ROI, cluster, hemisphere and stimulus type

To evaluate the response profiles of the ROIs and clusters (in both hemispheres) at a finer grain, and irrespective of whether there is selectivity for any of the text conditions over other categories, we attempted to decode, pairwise, the text condition from the multivariate pattern in each ROI and then in each cluster (i.e., decoding accuracy for words against letter strings, inverted words against letter strings, and words against inverted words) (Figure 5). We also derived the decoding accuracy (at the level of the ROI and cluster) for words, inverted words and letter strings, against the objects condition (see Appendix B, Figure 8).

**Figure 5:**
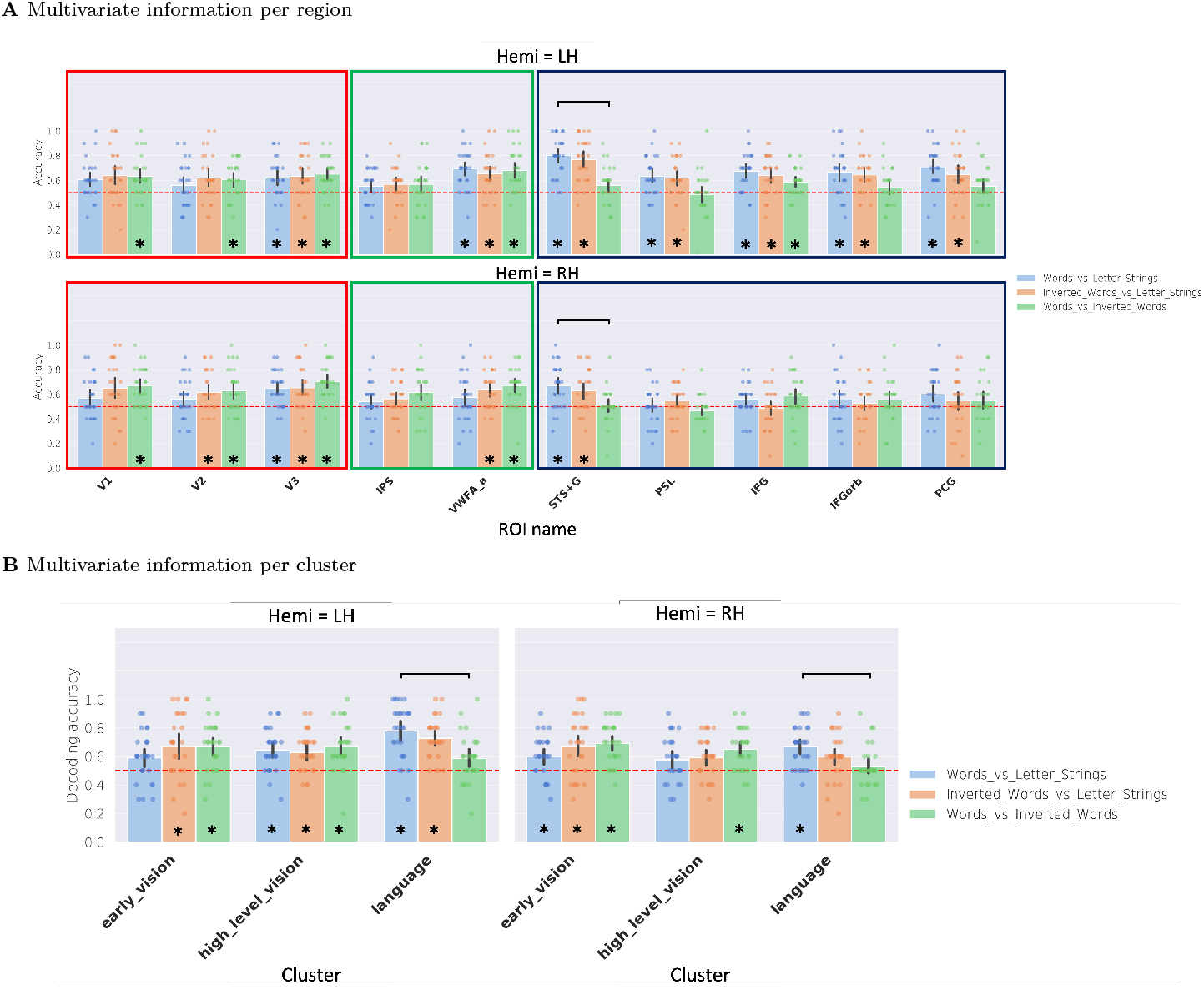
Multivariate decoding between the three text stimuli. Bars with asterisks are significantly greater than chance, computed using a one-sample *t*-test against 0.5 and corrected for multiple comparisons. **A**. Multivariate decoding accuracy for each pair of text conditions (i.e., Words versus Letter Strings, Inverted Words versus Letter Strings and Words versus Inverted Words) in each ROI of each hemisphere. Black lines indicate all significant within-hemisphere and within-cluster pairwise differences across stimulus types (horizontal lines). No between-ROI (within the same cluster) or between-hemisphere pairwise differences (across homologous ROI-stimulus combinations) survived post-hoc correction. **B**. Multivariate decoding accuracy within each of the three clusters in each hemisphere. Horizontal black lines indicate all significant within-hemisphere pairwise differences (across stimulus types). None of the between-cluster (within the same hemisphere) comparisons or between-hemisphere comparisons survived correction. In each plot, dots show scores for individual subjects, bars show the average across subjects, and black error lines indicate the 95% confidence interval of scores across subjects. Significance of all pairwise differences were computed with Tukey’s Honest Significant Difference (HSD) test and *p* < 0.01.

A repeated measures ANOVA with decoding accuracy as the dependent measure and ROI, hemisphere and textual stimulus pair (henceforth referred to as stimulus pair) as within-subject factors did not reveal a statistically significant three-way interaction (F(18, 486) = 0.9, *p* = 0.51, *η*^2^ = 0.03). However, all two-way interactions were significant: ROI×Hemisphere (F(9, 243) = 4.9, *p* < 0.001, *η*^2^ = 0.16); ROI×Stimulus-pair (F(18, 486) = 6.2, *p* < 0.001, *η*^2^ = 0.19); and Hemisphere × Stimulus-pair (F(2, 54) = 15.8, *p* < 0.001, *η*^2^ = 0.37). Two of the three main effects were also significant: ROI (F(9,243) = 6.48, *p* < 0.001, *η*^2^ = 0.19), and Hemisphere (F(1, 27) = 11.8, *p* < 0.005, *η*^2^ = 0.3). The main effect of Stimulus-pair was not significant (F(2, 54) = 2.2, *p* > 0.10, *η*^2^ = 0.07).

All ROIs that exhibited a significantly greater decoding accuracy than chance (0.5) for specific stimulus-pairs, as well as all pairwise comparisons within each of the hemispheres that survived post-hoc Tukey correction, are shown in Figure 5A. In the LH, all ROIs except the IPS were able to decode at least one pair of stimulus types with an accuracy significantly above chance. All Early vision ROIs could decode Words from Inverted Words, while all Language ROIs decoded Words and Inverted Words from Letter Strings, but most could not distinguish between Words and Inverted Words. The LH VWFA exhibited significant decoding accuracy for all three stimulus-pairs.

In contrast, in the RH, V1, V2, V3, VWFA and STS+G were the only ROIs whose decoding accuracy was significantly greater than chance (for at least one stimulus-pair). Similar to the LH, the voxel activation pattern in V1, V2, V3 and VWFA, but not STS+G, could decode Words from Inverted Words.

To assess decoding accuracy at the level of the cluster, we repeated the same ANOVA as above but with Cluster as a factor rather than ROI. There was neither a statistically significant three-way interaction (F(4, 108) = 0.4, *p* = 0.8, *η*^2^ = 0.01), nor a significant two-way interaction of Stimulus-pair × Hemisphere (F(2, 54) = 2.2, *p* = 0.1, *η*^2^ = 0.07). However, the two-way interactions of Cluster × Hemisphere (F(2, 54) = 8.0, *p* < 0.001, *η*^2^ = 0.23) and Cluster × Stimulus-pair (F(4, 108) = 13.0, *p* < 0.001, *η*^2^ = 0.33) were significant. There was also a significant main effect of Hemisphere (F(1, 27) = 12.4, *p* < 0.005, *η*^2^ = 0.32), but not of Stimulus-pair (F(2, 54) = 0.3, *p* > 0.50, *η*^2^ = 0.01) or Cluster (F(2, 54) = 0.6, *p* > 0.50, *η*^2^ = 0.02).

Figure 5B represents the specific stimulus-pairs for which each of the clusters exhibited a significantly high decoding accuracy (i.e., significantly greater than chance), and also pairwise comparisons within each hemisphere that survived post-hoc Tukey correction. As evident, the Early vision and High-level vision clusters in both the LH and RH could decode Words from Inverted Words. This is in contrast to the Language cluster in both hemispheres, which exhibited significant decoding accuracy for Words and Inverted Words against Letter Strings, but failed to decode Words from Inverted Words. None of the between-hemisphere pairwise comparisons (across homologous cluster-stimulus combinations) survived post-hoc correction, suggesting a very similar pattern of decoding accuracy across the three clusters in both hemispheres (for all three stimulus types), in strong contrast to the pattern found for selectivity.

In summary, evaluating decoding accuracy provided information over and above that gleaned from the univariate analysis. Specifically, we observed similar overall decoding accuracy in the LH and RH (both for decoding textual stimuli from each other, and textual stimuli against objects), even though selectivity of the RH was significantly lower than that of the LH for ROIs as well as clusters (Figures 4A and 4B). As evident in Figure 5, in both hemispheres, the accuracy of decoding words from inverted words decreased from the posterior (primarily ROIs in the Early vision cluster) to more anterior regions (ROIs in the Language cluster). The only notable exception to this trend was the VWFA, which, despite being more anterior to the ROIs of the Early vision cluster, exhibited similar, if not greater, decoding accuracy for words against inverted words compared to early visual regions (along with a similarly significant decoding accuracy for the other two stimulus-pairs). A similar posterior-to-anterior transition was seen at the cluster-level, such that the Early vision cluster had the highest decoding accuracy for words against inverted words, but the lowest decoding accuracy for words against letter strings, in both hemispheres. This pattern was reversed in the Language cluster, which displayed significantly higher decoding accuracy for words versus letter strings compared to words versus inverted words. This was likely primarily driven by the STS+G, which was the only ROI in the Language cluster to display significant differences in decoding accuracy across stimulus-pairs. Combined with the selectivity analyses, the strong multivariate information for distinguishing words and inverted words from letter strings in the STS+G provides converging evidence for a role of the RH STS+G in visual word processing.

### Functional connectivity of the visual word processing network

The focus, thus far, has been on individual ROIs and the aggregation of these regions into clusters. Here, we examine the interrelationships among ROIs and among clusters by documenting their functional connectivity profiles, and the modulation of these interrelationships by stimulus type.

We derived the FC of all ROI pairs within each hemisphere, as well as all homologous ROI pairs across hemispheres. Because the within-LH, within-RH and between hemispheres could not be captured in an orthogonal factorial analysis, we performed three separate ANOVAs, one for all ROI pairs within the LH (*‘LH-comparisons’*), one for the ROI pairs within the RH (*‘RH-comparisons’*) and, the last between all homologous (but not non-homologous) ROI pairs in the LH and RH (*‘between-hemisphere’*). For the factorial analysis of variance, we refer to the first ROI in a pair as ROI1, and the second as ROI2. As the partial functional correlation is symmetric, the pair index label is arbitrary; thus, where the effect of interest involves only a single ROI as in the cross-hemisphere homotopic regions (e.g., ROI × hemisphere), we do not use the ROI pair index label. Below, we present the heatmaps of words against the null distribution and then the partial functional correlation between ROI pairs for words compared to inverted words and letter strings.

For the *LH-comparisons* analysis, with ROI1, ROI2 and stimulus type as within-subject factors and FC as the dependent measure, we observed a significant three-way interaction (F(162,4374) = 2.07, *p* < 0.001, *η*^2^ = 0.07). The two-way interaction of ROI1 × ROI2 (F(81,2187) = 720.41, *p* < 0.001, *η*^2^ = 0.97) was also significant, but not ROI1/ROI2 × Stimulus (F(18,486) = 0.98, *p* = 0.48, *η*^2^ = 0.03). There were also significant main effects of ROI (F(9,243) = 14.9, *p* < 0.001, *η*^2^ = 0.36), but not of Stimulus (F(2,54) = 1.51, *p* = 0.23, *η*^2^ = 0.05). IFG was the ROI with the largest overall partial functional correlation with other regions, followed by STS+G, VWFA and then IFGorb.

As with the *LH-comparisons* analysis, we conducted the *RH-comparisons* analysis with ROI1, ROI2 and stimulus type as within-subject factors, and FC as the dependent measure. We observed a significant three-way interaction (F(162,4374) = 1.26, *p* = 0.01, *η*^2^ = 0.04). There was also a two-way interaction of ROI1 × ROI2 (F(81,2187) = 751.3, *p* < 0.001, *η*^2^ = 0.97), but not ROI × Stimulus (F(18,486) = 1.06, *p* = 0.38, *η*^2^ = 0.04). There was also a significant main effect of ROI (F(9,243) = 13.9, *p* < 0.001, *η*^2^ = 0.34), but not of Stimulus (F(2,54) = 0.51, *p* = 0.60, *η*^2^ = 0.02). IFG was the region most strongly connected to others followed by V2 and then STS+G.

To identify the ROI pairs that were significantly functionally correlated in the words stimulus condition, we computed one-sample post-hoc *t*-tests of the partial functional correlation of all ROI pairs in the LH and RH for words against a null distribution (leftmost panel; Figures 6A and 6B). To further illustrate the findings of the above-described ANOVAs and highlight the key interrelationships (between ROI pairs and across stimulus types), we also computed two-sample *t*-tests comparing the partial functional correlation of ROI pairs for words with that for letter strings and inverted words (Figures 6A and 6B; middle panel for letter strings, rightmost panel for inverted words). Asterisks denote ROI pairs that survived post-hoc Bonferroni correction (*p* < 0.0005). As evident from the heatmaps, there is considerable similarity across stimulus types, with significant connectivity between V1, V2 and V3, between the IPS and VWFA, and also between STS+G, PSL, IFG, IFGorb and PCG, albeit all to a slightly stronger degree in LH than RH. Although it is not surprising to see this connectivity pattern for words (as the clustering in Figure 3 was derived from the pairwise correlations of activations in ROIs in response to words), for the most part, similar pairwise correlations are seen across all stimulus types. Notably, the only significant stimulus difference that was seen was in the FC between VWFA and V3 comparing words and inverted words, with this connection being weaker under word inversion.

**Figure 6:**
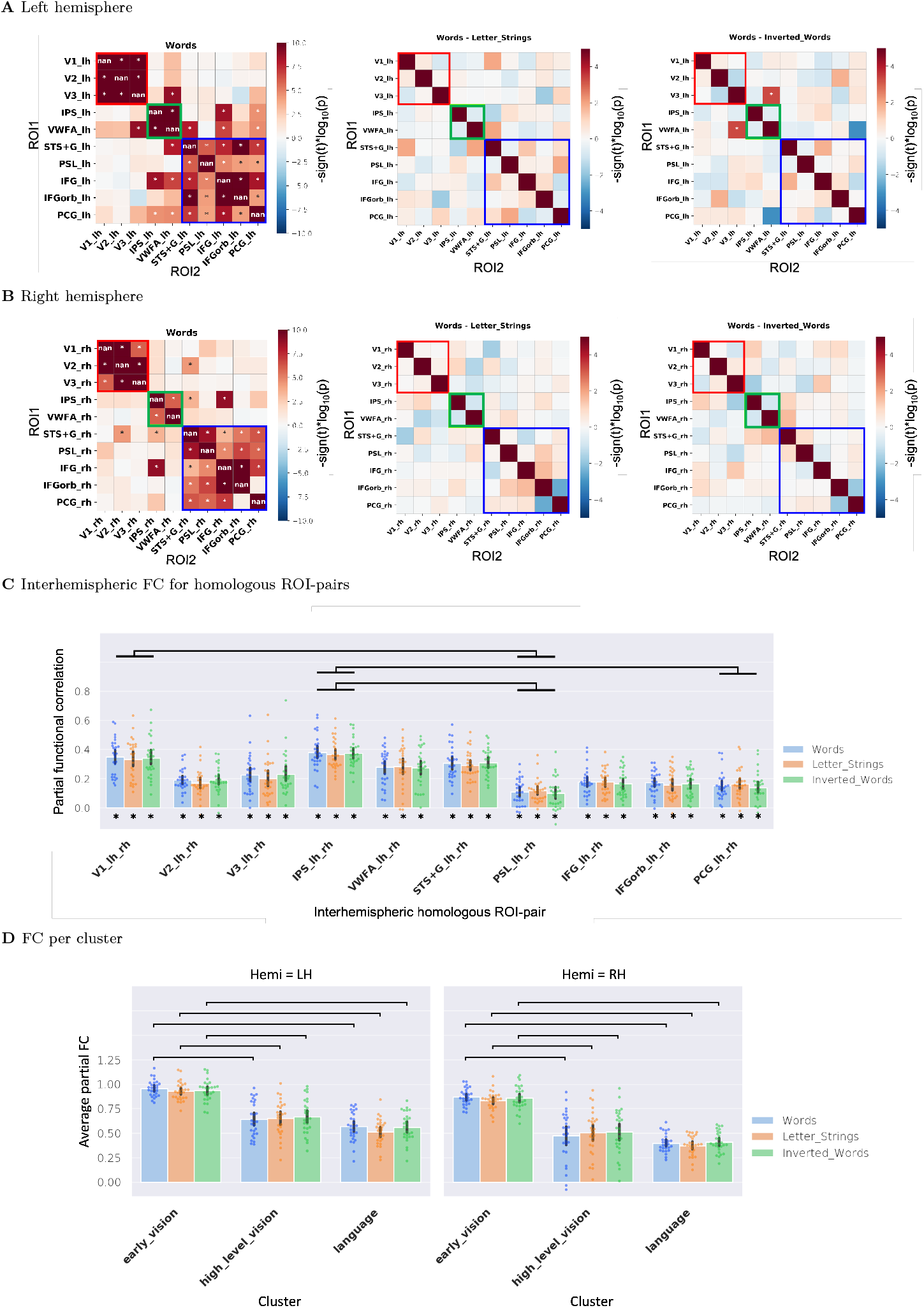
Functional correlation (connectivity) of all ROI pairs in the left hemisphere (**A**) and the right hemisphere (**B**) for the three stimulus conditions. The leftmost panel in **A** and **B** is a one-sample *t*-test of the partial functional correlation between ROI pairs for Words against the null distribution. The middle and rightmost panels in **A** and **B** are two-tailed *t*-tests of the partial functional correlation between ROI pairs for Words compared to Letter Strings and Inverted Words, respectively. Solid red, green and blue boxes surround ROI pairs in the Early vision, High-level vision and Language clusters respectively. Cells with asterisks represent partial functional correlations that survived Bonferroni correction (*p* < 0.0005). **C**. Partial FC of each of the interhemispheric homologous ROI-pairs for each of the three stimulus types. Horizontal black lines indicate all ROI-pairs that have significantly different interhemispheric partial FC, regardless of stimulus type. Bars with asterisks are significantly greater than 0, computed using a one-sample *t*-test against the null distribution (i.e., 0) and corrected for multiple comparisons. **D**. Average within-cluster partial FC for each of the three stimulus types in both hemispheres. Horizontal black lines indicate the top 70% of all significant within-hemisphere pairwise differences (across clusters and stimulus types). None of the between-hemisphere comparisons survived post-hoc correction. In both **C** and **D**, dots show scores for individual subjects, bars show the average across subjects, and black error lines indicate the 95% confidence interval of scores across subjects. Significance of all pairwise differences were computed with Tukey’s Honest Significant Difference (HSD) test and *p* < 0.01.

The *between-hemisphere* analysis revealed no interaction between ROI pair (i.e., pairs of homologous ROIs, for example, the FC value for LH-RH VWFA, labelled ‘VWFA pair’) and stimulus type (F(18,486) = 1.3, *p* > 0.1, *η*^2^= 0.05), nor a main effect of stimulus (F(2,54) = 1.8, *p* > 0.1, *η*^2^ = 0.06). There was, however, an effect of ROI pair (F(9,243) = 22.7, *p* < 0.001, *η*^2^ = 0.46). A one-sample *t*-test revealed that all ten between-hemisphere ROI pairs had significant partial functional correlation compared to the null distribution for all three stimulus types, and corrected for multiple comparisons (*p* < 0.001). The IPS emerged as having the strongest interhemispheric partial FC, followed by V1 and STS+G, based on a post-hoc Tukey HSD test (Figure 6C). These results highlight the existence of significant interhemispheric communication in ROIs of each of the three network clusters, regardless of stimulus type.

We also determined whether certain clusters showed higher FC than others and whether this differed as a function of stimulus type. Note that we used FC initially to calculate the clustering based just on the words condition (see Figure 3) so the analysis of the activation for words is not independent from the definition of clusters; however, this clustering was performed on group-averaged FC whereas analyses here are performed within individuals, and this does not affect the other conditions. In any case, caution should be taken when comparing the strength of word FC within versus between clusters. In a three-way ANOVA with cluster, hemisphere and stimulus type as within-subject factors, there was neither a significant three-way interaction (F(4, 108) = 1.1, *p* > 0.3, *η*^2^ = 0.04), nor two-way interactions of either Hemisphere × Cluster (F(2, 54) = 2.6, *p* > 0.05, *η*^2^ = 0.09) or Hemisphere × Stimulus (F(2, 54) = 1.3, *p* > 0.2, *η*^2^ = 0.05). The only significant interaction was between Cluster × Stimulus (F(4, 108) = 5.1, *p* < 0.001, *η*^2^ = 0.16). For all three stimulus types, there was greater FC for the Early vision, then High-level vision and, last, Language clusters, especially for words, and there was no difference for letter strings or inverted words. All of these comparisons survived correction using Tukey’s HSD test and *p* < 0.01 (Figure 6D; for the full list of all significant cluster functional connectivity comparisons that survived the post-hoc Tukey HSD correction, see Appendix C, Table 7). While the main effect of Stimulus was not significant (F(2, 54) = 2.1, *p* > 0.10, *η*^2^ = 0.07), there was a main effect of Hemisphere (F(1, 27) = 43.1, *p* < 0.001, *η*^2^ = 0.62), reflecting the greater FC in the LH than RH, and of Cluster (F(2, 54) = 102.8, *p* < 0.001, *η*^2^ = 0.79), with significantly greater FC in Early vision, followed by High-level vision, and with lowest FC in the Language cluster.

In summary, FC was highly modulated by hemisphere and ROI pair, but less by stimulus type. Pairwise connectivity was strong between V1-V3, between IPS and VWFA and between the remaining language regions, although almost always to a greater degree in the LH than RH. The LH VWFA exhibited significant FC with the largest number of ROIs, spanning across all three clusters. While some ROI pairs (both within and between hemispheres) exhibited a small difference in partial functional correlation for inverted words and letter strings compared to words, few comparisons survived post-hoc correction. All ROIs exhibited significant interhemispheric connectivity to their contralateral homologue, for all three stimulus types. The connectivity at the cluster level demonstrated greater LH than RH connectivity and stronger connectivity of the Early vision, then High-level vision clusters followed by the Language cluster for all three stimulus types. Although this reliable pattern of FC across stimulus types (at both the level of the ROI-pair and cluster) may seem surprising, we confirmed this pattern using only the peak of the hemodynamic response function (HRF) for the middle of each stimulus block (see Appendix C, Figure 10) and using the beta coefficients associated with each stimulus block obtained from the GLM, across all subjects (see Appendix C, Figure 11). Hence, it appears that although the activity of individual ROIs (in their univariate selectivity and multivariate decoding accuracy) is modulated by stimulus type, the interactions between pairs of ROIs, as well as average FC of the three clusters of ROIs remains largely stable across words, inverted words and letter strings.

### Node strength across stimulus types, and hemispheric differences

Following the comparisons of individual ROIs/clusters against each other and then pairs of ROIs/clusters against each other, in this final set of analyses, we evaluated the organization of the entire bihemispheric word processing circuit. To do so, we adopted a graph-theoretic approach and calculated node strength of each ROI, and then, separately, node-cluster strength of each ROI (defined below).

For a given ROI, node strength was computed as the total sum of the absolute partial functional correlation between that ROI and all other ipsilateral ROIs as well as its contralateral homologue (i.e., the sum of the absolute weights of all edges attached to that ROI, other than the edges between the ROI and non-homologous contralateral ROIs). The contralateral homologue was included as our FC analyses revealed significant interhemispheric homologous functional connectivity for all ROIs (Figure 6C). Moreover, previous research has provided evidence of significant structural connectivity between contralateral homologous ROIs through the corpus callosum. Connectivity between non-homologous ROIs across hemispheres is less clear.

The node-cluster strength for a given ROI and a given cluster was calculated as the sum of the absolute partial functional correlation between the ROI and all other ipsilateral ROIs within the cluster (in this case, summation was restricted to only the *ipsilateral* edges between the given ROI and cluster).

A repeated-measures ANOVA with ROI, hemisphere and stimulus type as within-subjects factors and node strength as the dependent measure revealed neither a significant three-way interaction (F(18, 486) = 0.71, *p* = 0.80, *η*^2^ = 0.03), nor significant two-way interactions of Stimulus × Hemisphere (F(2, 54) = 2.05, *p* = 0.14, *η*^2^ = 0.07) or ROI × Stimulus (F(18, 486) = 1.30, *p* = 0.18, *η*^2^ = 0.05). However, the two-way interaction of ROI × Hemisphere (F(9, 243) = 8.59, *p* < 0.001, *η*^2^ = 0.24) was significant. Figure 7A shows the node strength of individual ROIs as a function of stimulus type and hemisphere. Post-hoc Tukey HSD testing revealed a host of significant differences in node strength as a function of ROI and hemisphere. As shown by the vertical black lines between the LH and RH graphs, node strength for most ROI-stimulus pairs was higher in the LH than RH. The horizontal black lines show the top 10% of all within-hemisphere pairwise differences. In the LH, the PSL had the lowest average node strength across all three stimulus types; thus, the strongest differences in node strength were between the PSL and most other ROIs (particularly V3, VWFA, STS+G and IFG), which were similar to each other. The STS+G exhibited the highest average node strength across all stimulus types, closely followed by IFG and VWFA. In the RH, IPS and IFG had higher average node strength compared to most other ROIs, but especially PSL and PCG. In contrast to the LH, the RH VWFA had the third lowest average node strength across stimulus types after PSL and PCG. In both hemispheres, none of the ROIs exhibited significant modulation of their node strength by stimulus type. For the full list of all significant node strength comparisons that survived the post-hoc Tukey HSD correction, see Appendix D.

**Figure 7:**
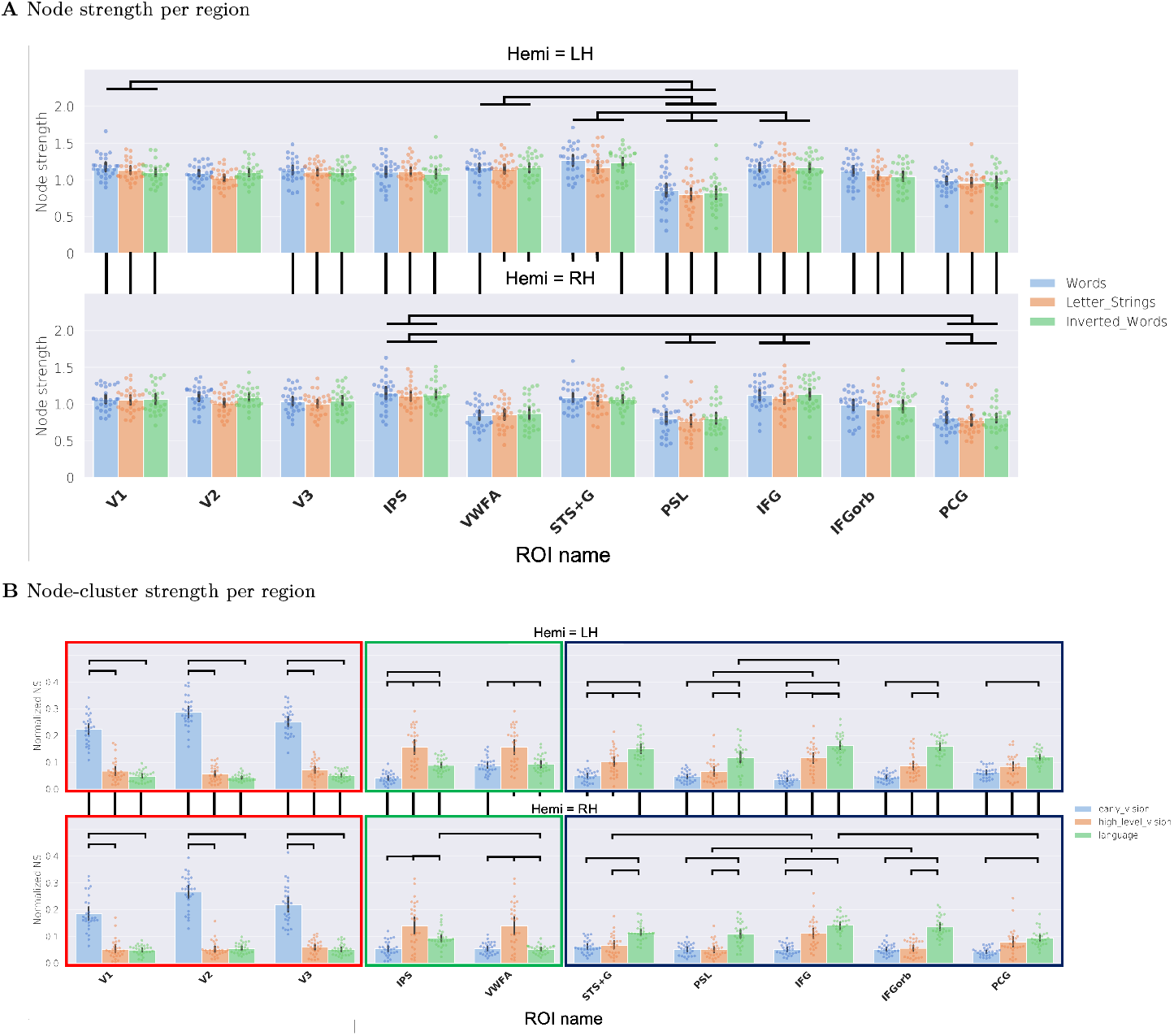
Effect of stimulus type and hemisphere on node strength (derived using partial FC) of each of the ROIs with the broad network of all other ipsilateral ROIs and the homologous contralateral ROI, as well as with each of the three clusters. **A**. The node strength of each ROI with the broader network of all other ROIs within the same hemisphere and the homologous ROI in the contralateral hemisphere, for each of the three stimulus types and across hemispheres. Horizontal black lines indicate the top 10% of all significant within-hemisphere pairwise differences, while the vertical lines indicate all between-hemisphere pairwise differences across homologous ROI-stimulus combinations. **B**. Normalized connectivity of each ROI to each of the three clusters (Early vision: blue bars, High-level vision: orange bars, and Language: green bars) in the LH (top panel) and the RH (bottom panel) for the Words stimulus condition. Vertical black lines indicate all significant between-hemisphere pairwise differences across homologous ROI-cluster combinations. Horizontal black lines indicate the top 30% of all within- and across-ROI comparisons in each hemisphere for the Early vision ROIs (surrounded by solid red box) as well as for the Language ROIs (surrounded by solid blue box). For the High-level vision ROIs (surrounded by solid green box), the horizontal lines represent the top 60% of all within- and across-ROI comparisons in each hemisphere (we show the most significant subset of comparisons here to avoid overcrowding the figure; a full list of pairwise comparisons can be found in Appendix D). In both **A** and **B**, dots show scores for individual subjects, bars show the average across subjects, and black error lines indicate the 95% confidence interval of the normalized node strength values across subjects. Significance of all pairwise differences was computed with Tukey’s Honest Significant Difference (HSD) test and *p* < 0.01.

All three main effects in the ANOVA were also significant: ROI (F(9, 243) = 27.5, *p* < 0.001, *η*^2^ = 0.50), Stimulus (F(2, 54) = 5.4, *p* < 0.01, *η*^2^ = 0.17), and Hemisphere (F(1, 27) = 34.6, *p* < 0.001, *η*^2^ = 0.56).

In summary, node strength, which measures, for each ROI, the strength of its connection with the broad cluster comprising all other ipsilateral ROIs and its contralateral homologue, is higher in the LH than RH across the board. Within each hemisphere, ROIs differed from each other in terms of the strength of their connection to the rest of the ROIs, but there was no significant modulation of the ROIs themselves by stimulus type. Hence, similar to the pairwise functional connectivity between ROIs, network-level analyses of the node strength of each ROI within the broader set of ROIs exhibits a stability across stimulus types.

### Differential node-cluster strength across ROIs, with high-level visual regions mediating between early vision and language

Finally, we assessed the connectivity of each of the ROIs to each of the three clusters: Early vision, High-level vision and Language, separately. This was computed as the sum of an ROI’s absolute partial functional correlation with all other ipsilateral ROIs in a cluster. This sum was then normalized by a factor *n−*1, where *n* is the total number of ROIs being considered (the subtracted 1 is to exclude self-connections). This computation is distinct from the average functional connectivity of the three clusters, which we have reported above (Figure 6D), as it focuses on the connectivity of nodes to clusters, rather than all connectivity within a cluster.

In Figure 7B, the boxes surrounding groups of individual ROIs are indicative of the cluster to which they belong: the Early vision cluster (red box), the High-level vision cluster (green box) and the Language cluster (blue box). Within each of these boxes, the y-axis reflects the normalized connectivity of the ROIs in a given hemisphere to their ‘*native*’ cluster (i.e., the cluster to which they belong, see Figure 3A) and with the other two ‘*non-native*’ clusters. For example, for the ROI V1, its native cluster would be Early vision, and its non-native clusters would be High-level vision and Language. For each ROI, the blue bar represents its connectivity to the Early vision cluster, the orange bar to the High-level vision cluster and the green bar to the Language cluster. Figure 7B depicts the node connectivity of each of the individual ROIs only for the words stimulus condition (figures for the letter strings and inverted words conditions are not significantly different, and are included in the Supplemental Material; see Appendix D, Figure 12).

A three-way repeated measures ANOVA with hemisphere, ROI and cluster as within-subject factors revealed a significant interaction of all three factors (F(18, 486) = 3.7, *p* < 0.001, *η*^2^ = 0.12). Detailed scrutiny of the data revealed the source of the three-way interaction. All ten ROIs (in both hemispheres) exhibited significantly greater normalized connectivity to their native cluster, compared to their non-native clusters. Moreover, an interesting pattern emerged across the ROIs in terms of their relative connectivity to their non-native clusters. The ROIs in the Early vision cluster (V1, V2 and V3; surrounded by the red box in Figure 7B) and the Language cluster (STS+G, PSL, IFG, IFGorb and PCG; surrounded by the blue box in Figure 7B) all shared the characteristic of having significantly higher connectivity to the High-level vision cluster than to their other non-native cluster (see Appendix D). This was true in both the LH and the RH. Among the ROIs of the High-level vision cluster (IPS and VWFA; surrounded by the green box in Figure 7B), the IPS exhibited significantly higher connectivity to the Language cluster compared to the Early vision cluster. On the contrary, unlike most other ROIs, the VWFA, in both the LH and the RH, exhibited very similar connectivity to its non-native clusters (Early vision and Language). Furthermore, VWFA had the highest connectivity to the Early vision and Language clusters compared to other ROIs for which either of those two clusters were non-native (this was also true in both hemispheres). Supplementing the findings shown in Figure 7A, most ROIs had significantly greater connectivity to the three clusters in the LH compared to the RH. For the full list of all significant node strength comparisons that survived the post-hoc Tukey HSD correction, see Appendix D.

The two-way interactions of ROI × Cluster (F(18, 486) = 143.9, *p* < 0.001, *η*^2^ = 0.84) and ROI × Hemisphere (F(9, 243) = 3.6, *p* < 0.001, *η*^2^ = 0.12) were also significant, but that of Cluster × Hemisphere was not (F(2, 54) = 1.0, *p* = 0.37, *η*^2^ = 0.04). Last, all three main effects were also significant: ROI (F(9,243) = 24.0, *p* < 0.001, *η*^2^ = 0.47), cluster (F(2,54) = 19.8, *p* < 0.001, *η*^2^ = 0.42), and hemisphere (F(1, 27) = 49.1, *p* < 0.001, *η*^2^ = 0.65).

To summarize, the results of our graph-theoretic analyses suggest that, bihemispherically, the three clusters (Early vision, High-level vision and Language) function as ‘subnetworks’ of a broader visual word processing network, as they show greater within-than between-cluster connectivity, and each of the ROIs shows preferential connectivity with their native cluster (this is true for inverted words and letter strings as well; see Appendix D, Figure 12). However, while a general preference for connectivity to native clusters is expected, and was generally confirmed, differential weighting of connectivity to the non-native clusters across ROIs emerged as well. All ROIs in the Early vision and Language clusters had higher connectivity to the High-level vision cluster than to their other non-native cluster. In contrast, the VWFA, in both the LH and RH, showed similar connectivity to both of its non-native clusters (Early vision and Language). This suggests that the High-level vision cluster, and in particular the VWFA, may play a role in mediating signal propagation and integration between the Early vision and Language clusters. Our collective univariate, multivariate and functional connectivity results further reinforce the theory that the VWFA may be the more dominant of the two ROIs in the High-level vision cluster in terms of its mediating function, as discussed in greater detail in the Discussion. Moreover, similar to the generic node strength of each of the ROIs within the broad set of ROIs (Figure 7A), the normalized connectivity of each ROI to the three clusters is also relatively stable and unmodulated across stimulus types.

## Discussion

The goal of this study was to characterize comprehensively in a largely data-driven fashion those regions in human cortex that show selective responses to orthographic input. Decades of studies have attested to the key role of the Visual Word Form Area (VWFA) in the LH in the task of word processing. This primary focus on the VWFA and its seemingly singular role in word processing has, however, been challenged by investigations showing activation to text beyond the VWFA including in Broca’s and Wernicke’s area as well as the Intraparietal Sulcus, Prefrontal Gyrus and Superior Temporal and Supramarginal gyri (Chen et al., 2019; Stevens et al., 2017).

The first aim of the present study was to define, within individual, the large-scale distributed circuit for word processing as well as the functional properties and interrelationships between the different regions and in both hemispheres. Therefore, based on whole-brain fMRI data obtained while participants viewed three different textual stimulus types (words, letter strings and inverted words) and non-text (faces, inverted faces and objects) conditions, we examined responses in a set of 10 regions in each hemisphere and at an intermediate level of organization derived from a clustering algorithm that grouped the ROIs into three clusters: Early vision, High-level vision, and Language. For each ROI and cluster in each hemisphere, we quantified the univariate response selectivity profile, the multivariate decoding accuracy, and the pairwise functional connectivity (FC). Last, using graph theory metrics, we measured node strength, with each ROI as a node within the broader network of all ten ROIs, and as a node within each of the three defined clusters.

The analysis of univariate ROI selectivity in the LH revealed significant selectivity in most ROIs for at least one of the three text stimulus types (see Figure 4), with ROIs of the Early vision cluster evincing similar selectivity for each of the text stimulus types, and the High-level vision cluster showing more selectivity for letter strings than the other conditions. The VWFA exhibited the highest average selectivity across text stimulus types of all the ROIs, replicating the standard profile reported ubiquitously in the literature. Last, most ROIs in the Language cluster exhibited significant selectivity to words and inverted words, and less to letter strings (see Figure 4). These findings were supplemented by multivariate results revealing that at least one pair of textual stimuli could be decoded in most LH ROIs (see Figure 5; see Appendix B, Figure 8) with all three pairs of textual stimuli decodable from each other in VWFA. Words and letter strings were better decoded than inverted words in the Early vision cluster suggesting representation of visual features (i.e., upright versus inverted text). The ROIs of the Language cluster had at-or below-chance decoding accuracy for words against inverted words, but exhibited very high accuracy for both stimuli against letter strings, emphasizing the lexical aspects of the input. All of these findings were replicated in the univariate and multivariate analyses when sampling from the three clusters in the LH with, again, similar significant modulation by stimulus type.

For the RH, selectivity was also identified in the ROI univariate and multivariate analyses. RH univariate selectivity for text was found consistently only in early visual regions and in the STS+G for words. Thus, intriguingly, even early visual regions seemed to demonstrate a leftward bias, implying that the observed selectivity was not merely a function of form properties differing between stimulus categories. While the right VWFA anatomical correlate did not contain significant univariate selectivity, multivariate analyses revealed significant decodability of text conditions from each other, similarly to RH STS+G. The early visual regions of the RH decoded textual stimuli similarly to the LH. Besides STS+G, no other language regions showed consistent text selectivity, nor consistent decodability of text conditions. These findings highlight the partial involvement of the RH in text processing, typically limited to more posterior regions. The STS+G in particular, seems to be activated by the lexical or semantic content of word-like stimuli, and may play an important role in feedback to other regions of the word processing network.

Analyses of pairs of ROIs revealed strong functional connectivity (FC) between ROIs within a cluster (i.e., V1-V3, between IPS and VWFA and between the remaining language regions), as expected, although not as strongly within the high-level vision cluster, which connected strongly with both Early vision and Language ROIs. A similar pattern was observed at the level of the cluster (see Figure 6). While greater within versus between cluster connectivity was strongly expected for the Words condition as the clustering solution was defined on the basis of the group averaged

Words FC, the same patterns of FC were also observed in the other conditions (Appendix D, Figure 12); moreover, the other conditions generated the same intermediate clustering solutions (see Appendix C, Figure 9). Thus, the greater within vs. between cluster connectivity cannot simply be explained in terms of double dipping (Kriegeskorte et al., 2008). The FC profile of the RH mirrored that of the LH, although slightly weaker, as reflected both in the *t*-test between ROI pairs for Words against the null distribution and in the two-tailed *t*-tests between ROI pairs for Words compared to Letter Strings and Inverted Words, respectively. The parallel arrangements of the LH and RH were also revealed in the FC interhemispheric analysis: homologous pairs of ROIs across hemispheres were more strongly connected than non-homologous pairs, with V1, IPS, VWFA and STS+G being the most strongly functionally connected to their contralateral homologues. Of particular interest, both within-and between-hemisphere FC between most pairs of ROIs held to an equivalent extent across stimulus types.

The final analysis explored circuit wide connectivity by calculating the node connectivity of each ROI with all other ROIs or with each of the three clusters separately. Node strength of most ROIs was consistently greater in the LH than RH, and similar to FC, exhibited little modulation by stimulus type (see Figure 6). In the LH, across all stimulus types, STS+G and VWFA exhibited the highest average node strength while PSL had the lowest average node strength but, in the RH, compared with most other ROIs, the VWFA had significantly lower node strength, and IFG, IPS and STS+G had higher node strength. In both hemispheres, most ROIs did not exhibit significant modulation of their node strength by stimulus type. Bihemispherically, the three clusters appeared to serve as ‘subnetworks’ of a broader visual word processing network, with greater within-than between-cluster connectivity, and with each of the ROIs exhibiting preferential connectivity to its native cluster. Of note, though, there was also differential weighting preferences of the non-native clusters for each ROI—all ROIs in the Early vision and Language clusters exhibited higher connectivity to the High-level vision cluster than to their other non-native cluster, implicating the High-level vision cluster in mediating information flow and integration between the Early vision and Language clusters. Unlike any of the other ROIs, the VWFA in both hemispheres showed highly similar node connectivity to its non-native clusters (Early vision and Language).

One noteworthy outcome of this study is the identification of several regions involved in visual word processing in the LH, with IFG, the Precentral Gyrus and the Superior Temporal Sulcus and Gyrus showing significant univariate activation (greater selectivity for words > inverted words > letter strings) and decoding accuracy almost as high as VWFA. That this result has not been reported widely is likely a function of the fact that most studies focus specifically on the VWFA. Along similar lines, several regions in the RH also exhibited significant selectivity and decoding accuracy. The RH STS+G showed the strongest sensitivity to text as evident in univariate selectivity, decoding accuracy and even FC profile, particularly compared to other RH regions such as IPS and VWFA. All ROIs displayed strong interhemispheric connectivity with their homologous contralateral counterpart. Hence, together, these various analyses bring out subtleties in the interactions among this broad network of ROIs that spans both hemispheres, as well as the selective modulation of this network by differing stimulus types (as described below).

It is interesting that many ROIs exhibit stimulus-dependent differences in their intrinsic selectivity and decoding ability among text conditions. However, when interactions between these ROIs are analyzed (i.e., their pairwise functional connectivity or their node strength within an inter-dependent interconnected cluster of ROIs), they are less modulated by stimulus type and, instead, exhibit stability. This is reflected in the largely invariant response to different stimuli within and largely across hemispheres, as well. This stimulus-independent profile of the interconnected cluster of ROIs confers a well-defined structure to the overall visual word processing network, which comprises widespread cortical areas associated with early and higher-level visual cortex, and language regions. On the other hand, the stimulus-dependent selectivity and multivariate voxel-wise pattern of the individual ROI entities, demonstrates the perceptual sensitivity of the network’s responses to stimulus types. The simultaneous presence of response flexibility and connectivity stability is consistent with artificial neural networks, in which different stimuli engage different representations to differing degrees even with a single set of weights, and with studies of physiology in which population level activity maintains consistent properties so that the network exhibits homeostasis (Driscoll et al., 2017, 2022).

We have found several indications of involvement of the RH, particularly the STS+G ROI. The obvious question, though, is whether RH engagement is epiphenomenal or plays a functional role. One possibility is that the RH has no involvement in word processing and that through tight coupling of homologous areas via the callosum, activation might merely cascade interhemipherically from the LH. If this were the case, then we might expect all RH regions to ‘inherit’ the activation from the LH but this is not the case at all and early visual regions, VWFA and STS+G are more prominently responsive than the other ROIs. Thus, a more parsimonious explanation is that the particular RH activation we discovered is functionally relevant.

Recent findings confirm that the RH homologue of Wernicke’s area (STS+G) retains some language capabilities even if as a ‘weak shadow’ to the LH prowess in the mature brain (Newport et al., 2022; Martin et al., 2022). These language capabilities appear to suffice for word processing following left occipitotemporal cortex resection, especially in childhood, with the RH VWFA activated by orthography in many cases (Cohen et al., 2004; Liu et al., prep). Relatedly, children with a single hemisphere (post-hemispherectomy for the management of medically resistant epilepsy) score, on average, 80 percent accuracy on a word discrimination task and this is so even with just a preserved RH (Granovetter et al., 2022). Also, in adult patients with prosopagnosia following focal damage to the RH, a moderate reading deficit, less severe than after LH damage, can be detected (Behrmann and Plaut, 2014) and, in an adult patient with (partially recovered) pure alexia, stimulation of the RH but not the LH disrupted oral reading (Coslett and Monsul, 1994). Together, these data also reveal meaningful and functional involvement of RH circuitry in word processing. Further investigation of the contribution of the RH to text processing is warranted, especially in the context of causal approaches (Planton et al., 2022).

Another outcome of this research is that, using a data-driven clustering approach, we found that the ROIs aggregated into clusters easily labeled as Early vision, High-level vision and Language. Moreover, within the LH High-level vision subnetwork, the well-situated VWFA appears to play a more prominent mediating role than IPS as evident from the consistently stronger results across all analytic approaches. The LH VWFA exhibits significantly greater selectivity compared to the IPS for all three stimulus types, can differentiate between the three textual stimulus types more accurately compared to the IPS and displays significant functional connectivity with the largest number of ROIs across all three clusters (Figure 6). These characteristics of the VWFA allow it to play a more ‘privileged’ role than IPS in mediating information between the Early vision subnetwork and the Language subnetwork. The geographic location of the LH VWFA is well-suited to its synthesis of bottom-up visuospatial input features from occipital cortex with more conceptual representations such as those involved in language processing (Price and Devlin, 2011). The hierarchical arrangement among the three subnetworks in the RH is less clear and the mediation role of the VWFA is present but less obvious presumably because of the RH’s ‘minor’, non-dominant language status.

## Conclusions and future directions

There has been a growing realization that regions of the brain are deeply interconnected, and that behavior is the emergent property of distributed networks rather than the outcome of computations in a single, circumscribed region of cortex (Tooley et al., 2022). Taken together, the strong evidence for engagement of the LH early visual, high-level vision and language areas, along with the provocative findings of the contribution of many of the same regions in the RH, suggest that the whole-brain organization of the word processing network is not simply confined to one hemisphere and to one region. Rather, multiple regions and both hemispheres are involved, exhibiting complex temporal and task-based dynamics (in terms of individual ROI flexibility and broad network stability). Despite bihemispheric engagement, the LH appears primary, consistent with increasing representational distinctiveness in the left VTC and increasing reading performance in children, beyond the effect of age (Nordt et al., 2022), and despite additional increases in pseudoword distinctiveness in the RH.

We have provided a detailed characterization of the structure (the multiplicity of nodes and edges) and behavior (stimulus-driven modulation) of the visual word processing network. Of course, our findings raise even more questions. The adoption of a task of single word sequential discrimination is unlike the fluent reading of the mature reader. Approximating a more naturalistic scenario of text reading, with its further complexity, is needed to advance a deeper understanding of the process of reading. A thorough behavioral assessment in parallel with elucidating the neural circuitry will inform a deeper understanding of hierarchical organization, and flexibility and stability associated with different task conditions (White et al., 2023). Moreover, temporal dynamics that unfold during the process of reading await further explication (Vida et al., 2017; Woolnough et al., 2022a), and individual and cultural differences both within- and between-hemispheres remain to be explained (Hirshorn and Harris, 2022; Kubota et al., 2022; Mao et al., 2021; Nordt et al., 2021; Shechter et al., 2022).

## Acknowledgement

This research was supported by grants from NSF to MB and DCP (BCS2123069), Undergraduate Research Fellowship to RV and a Neuroscience Institute Presidential Fellowship to NB. MB acknowledges support from P30 CORE award EY08098 from the National Eye Institute, NIH, and unrestricted supporting funds from The Research to Prevent Blindness Inc, NY, and the Eye & Ear Foundation of Pittsburgh. We are also grateful to Dr. Anne Margarette Maallo and Roshni Nischal for their assistance with data collection.

