## Supplementary material for "Beyond the Visual Word Form Area: Characterizing a hierarchical, distributed and bilateral network for visual word processing"

### Appendix

### A Selectivity

#### A.1 ROI-level

**Table 2:** Significant comparisons that survived post-hoc Tukey correction

| S. No. | ROI 1 | Stimulus 1 | Hemi 1 | ROI 2 | Stimulus 2 | Hemi 2 | Mean Difference |
|---|---|---|---|---|---|---|---|
| 1 | psl | w | lh | stsg | w | lh | -542.569 |
| 2 | pcg | w | lh | stsg | w | lh | -522.937 |
| 3 | ips | ls | lh | vwfa | ls | lh | -504.694 |
| 4 | ifgorb | w | lh | stsg | w | lh | -502.096 |
| 5 | ifg | w | lh | stsg | w | lh | -480.543 |
| 6 | ips | iw | lh | vwfa | iw | lh | -477.377 |
| 7 | psl | iw | lh | stsg | iw | lh | -474.145 |
| 8 | vwfa | ls | lh | vwfa | ls | rh | 468.819 |
| 9 | vwfa | iw | lh | vwfa | iw | rh | 455.349 |
| 10 | stsg | ls | lh | stsg | w | lh | -427.975 |
| 11 | ifgorb | iw | lh | stsg | iw | lh | -424.217 |
| 12 | pcg | iw | lh | stsg | iw | lh | -411.199 |
| 13 | stsg | w | lh | stsg | w | rh | 385.494 |
| 14 | stsg | iw | lh | stsg | iw | rh | 377.122 |
| 15 | ifg | iw | lh | stsg | iw | lh | -375.119 |
| 16 | ips | w | lh | vwfa | w | lh | -344.358 |
| 17 | stsg | iw | lh | stsg | ls | lh | 342.077 |
| 18 | vwfa | w | lh | vwfa | w | rh | 328.268 |
| 19 | psl | w | rh | stsg | w | rh | -242.921 |
| 20 | v1 | ls | lh | v1 | ls | rh | 235.43 |
| 21 | ifg | w | rh | stsg | w | rh | -232.363 |
| 22 | pcg | w | rh | stsg | w | rh | -224.003 |
| 23 | ifgorb | w | rh | stsg | w | rh | -221.681 |
| 24 | stsg | ls | rh | stsg | w | rh | -194.024 |
| 25 | psl | ls | lh | stsg | ls | lh | -183.711 |
| 26 | vwfa | iw | lh | vwfa | w | lh | 182.757 |
| 27 | vwfa | ls | lh | vwfa | w | lh | 176.016 |
| 28 | psl | iw | rh | stsg | iw | rh | -164.118 |
| 29 | pcg | ls | lh | stsg | ls | lh | -161.987 |
| 30 | v3 | ls | lh | v3 | ls | rh | 151.638 |
| 31 | stsg | ls | lh | stsg | ls | rh | 151.543 |
| 32 | v1 | w | lh | v1 | w | rh | 150.225 |
| 33 | v1 | ls | lh | v2 | ls | lh | 147.3 |
| 34 | pcg | iw | rh | stsg | iw | rh | -146.883 |
| 35 | ifgorb | ls | lh | stsg | ls | lh | -145.719 |
| 36 | ifgorb | iw | rh | stsg | iw | rh | -144.245 |
| 37 | ifg | w | lh | ifg | w | rh | 137.314 |
| 38 | v1 | ls | lh | v3 | ls | lh | 127.767 |
| 39 | ifg | iw | lh | ifg | iw | rh | 124.775 |
| 40 | ifg | iw | rh | stsg | iw | rh | -122.772 |
| 41 | v2 | ls | lh | v2 | ls | rh | 120.796 |
| 42 | stsg | iw | rh | stsg | ls | rh | 116.498 |
| 43 | v1 | ls | lh | v1 | w | lh | 113.546 |
| 44 | pcg | iw | lh | pcg | iw | rh | 112.806 |
| 45 | v1 | iw | lh | v1 | ls | lh | -110.494 |
| 46 | v1 | iw | lh | v1 | iw | rh | 109.032 |
| 47 | v1 | iw | lh | v3 | iw | lh | 108.22 |

| | | | | | | | |
|---|---|---|---|---|---|---|---|
| 48 | ifgorb | w | lh | ifgorb | w | rh | 105.079 |
| 49 | ifg | ls | lh | stsg | ls | lh | -102.596 |
| 50 | ifg | iw | lh | psl | iw | lh | 99.025 |
| 51 | ifgorb | iw | lh | ifgorb | iw | rh | 97.151 |
| 52 | v2 | iw | lh | v2 | iw | rh | 93.735 |
| 53 | pcg | iw | lh | pcg | ls | lh | 92.865 |
| 54 | v3 | iw | lh | v3 | ls | lh | -90.946 |
| 55 | v1 | w | lh | v3 | w | lh | 89.569 |
| 56 | v1 | iw | lh | v2 | iw | lh | 89.565 |

#### A.2 Cluster-level

**Table 3:** Significant comparisons that survived post-hoc Tukey correction

| S. No. | Cluster 1 | Stim 1 | Hemi 1 | Cluster 2 | Stim 2 | Hemi 2 | Mean Difference |
|---|---|---|---|---|---|---|---|
| 1 | lang | w | lh | lang | w | rh | 800.29 |
| 2 | lang | iw | lh | lang | iw | rh | 778.949 |
| 3 | lang | ls | lh | lang | w | lh | -685.741 |
| 4 | hv | w | lh | lang | w | lh | -682.933 |
| 5 | lang | iw | lh | lang | ls | lh | 619.719 |
| 6 | ev | w | lh | lang | w | lh | -512.702 |
| 7 | ev | ls | lh | ev | ls | rh | 507.864 |
| 8 | ev | iw | lh | lang | iw | lh | -473.677 |
| 9 | hv | ls | lh | hv | ls | rh | 462.787 |
| 10 | hv | iw | lh | hv | iw | rh | 446.235 |
| 11 | ev | ls | lh | lang | ls | lh | 400.242 |
| 12 | hv | iw | lh | lang | iw | lh | -384.416 |
| 13 | hv | w | lh | hv | w | rh | 350.038 |
| 14 | lang | ls | lh | lang | ls | rh | 325.165 |
| 15 | ev | w | lh | ev | w | rh | 292.656 |
| 16 | ev | iw | lh | ev | iw | rh | 267.848 |
| 17 | ev | iw | lh | ev | ls | lh | -254.199 |
| 18 | hv | w | rh | lang | w | rh | -232.681 |
| 19 | hv | iw | lh | hv | w | lh | 232.495 |
| 20 | ev | w | rh | hv | w | rh | 227.613 |
| 21 | ev | ls | lh | ev | w | lh | 227.202 |
| 22 | ev | ls | rh | lang | ls | rh | 217.543 |

### B MVPA

#### B.1 Decoding accuracy for textual stimuli against objects baseline

As displayed in Figure 8A, decoding accuracy was significantly above chance (0.5) for every ROI and stimulus type. A repeated measures ANOVA with decoding accuracy as the dependent measure and ROI, hemisphere and stimulus type as within-subject factors revealed a statistically significant three-way interaction (F(18, 486) = 3.2, $p < 0.001$, $\eta^2 = 0.11$) and all two-way interactions were significant: ROI × Hemisphere (F(9, 243) = 3.1, $p < 0.005$, $\eta^2 = 0.10$); ROI × Stimulus (F(18, 486) = 4.3, $p < 0.001$, $\eta^2 = 0.137$); and Hemisphere × Stimulus (F(2, 54) = 3.9, $p < 0.03$, $\eta^2 = 0.13$). The three main effects were also significant: ROI (F(9,243) = 68.6, $p < 0.001$, $\eta^2 = 0.72$), Hemisphere (F(1, 27) = 35.4, $p < 0.001$, $\eta^2 = 0.567$) and Stimulus (F(2, 54) = 3.9, $p < 0.03$, $\eta^2 = 0.13$).

Figure 8A displays the top 70% of all significant within-hemisphere pairwise comparisons (horizontal black lines) and between-hemisphere pairwise comparisons (vertical black lines). In general, most language-related regions exhibited higher decoding accuracy in the LH than RH for at least one of the three stimulus types. In the LH, there were no differences across stimulus types within or between ROIs for V1-V3. The VWFA had the highest decoding accuracy across all ROIs for all three stimulus types. Among the language-related regions, STS+G had the best decoding accuracy (particularly for words and inverted words), and PCG showed the most prominent modulation of decoding accuracy by stimulus type (higher for words and inverted words compared to letter strings). The RH exhibited a similar set of significant pairwise comparisons, except that STS+G, but not PCG, was significantly modulated by stimulus type.

A repeated measures ANOVA with decoding accuracy as the dependent measure and cluster, stimulus type and hemisphere as within-subject factors (see Figure 8B) indicated no significant three-way interaction (F(4, 108) = 0.50, $p = 0.74$, $\eta^2 = 0.018$). The two-way interactions of Stimulus × Cluster (greater decoding accuracy for words and inverted words compared to letter strings; F(4, 108) = 5.49, $p < 0.001$, $\eta^2 = 0.17$) and Cluster × Hemisphere (greater decoding accuracy of each of the clusters in the LH than RH; F(2, 54) = 5.01, $p < 0.05$, $\eta^2 = 0.16$) were significant, but Stimulus × Hemisphere was not significant (F(2, 54) = 0.17, $p = 0.84$, $\eta^2 = 0.006$). Last, there was a main effect of Cluster (F(2, 54) = 46.28, $p < 0.001$, $\eta^2 = 0.63$), with the highest decoding accuracy in the High-level vision cluster, followed by the Early vision and then Language clusters. There was also a main effect of Hemisphere, with greater decoding accuracy in the LH than RH (F(1, 27) = 41.86, $p < 0.001$, $\eta^2 = 0.61$). The main effect of Stimulus was not significant (F(2, 54) = 2.78, $p = 0.07$, $\eta^2 = 0.09$).

Post-hoc Tukey HSD with $p < 0.01$ revealed that, within the LH, decoding was more accurate in the High-level vision cluster compared to the Language cluster for all three stimulus types. The Early vision cluster exhibited greater decoding accuracy than the Language cluster only for the letter strings condition. Within the RH, decoding accuracy was higher in the Early and High-level vision clusters than in the Language cluster for all three stimulus types. None of the between-hemisphere comparisons (between homologous stimulus-cluster pairs) survived the Tukey HSD correction.

**A** Multivariate information per region

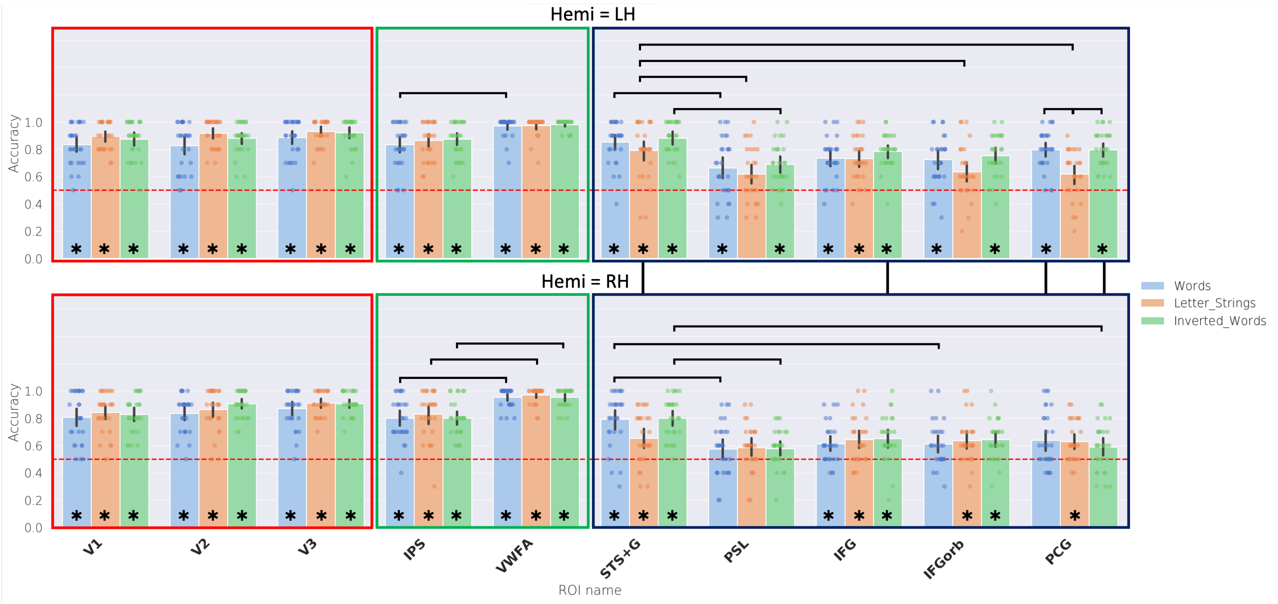


**B** Multivariate information per cluster

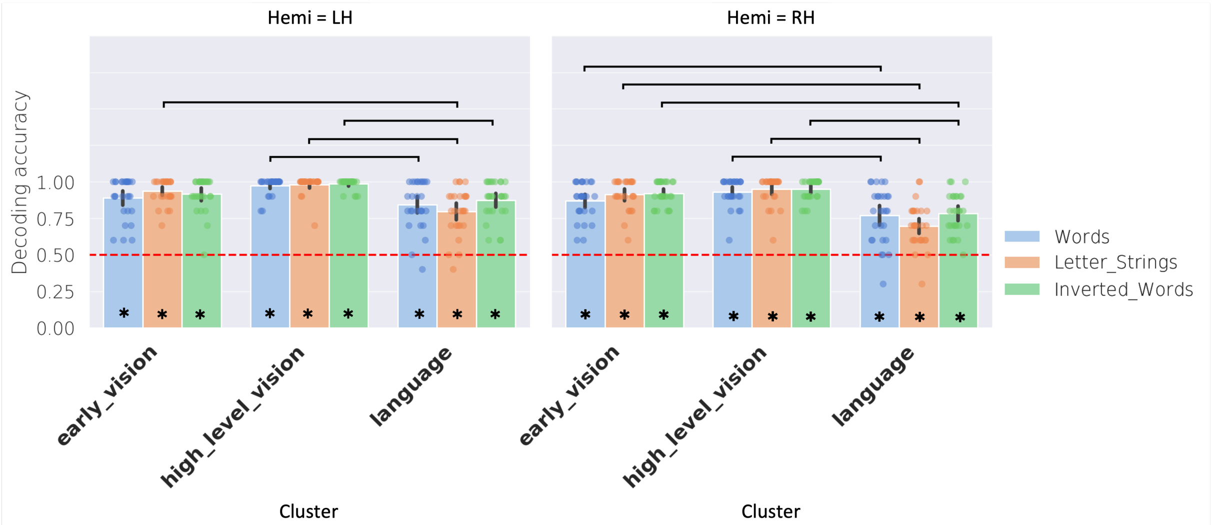


**Figure 8:** Multivariate decoding for text stimuli. **A.** Multivariate decoding accuracy for each text condition vs. objects in each ROI of each hemisphere, grouped by cluster. Black lines indicate the top 70% of all significant within-hemisphere pairwise differences across ROIs and stimulus types (horizontal lines) and between-hemisphere pairwise differences across homologous ROI-stimulus combinations (vertical lines). **B.** Multivariate decoding accuracy within each of the three clusters in each hemisphere. Horizontal black lines indicate all significant within-hemisphere pairwise differences (across clusters and stimulus types). None of the between-hemisphere comparisons survived correction. In each plot, dots show scores for individual subjects, bars show the average across subjects, and black error lines indicate the 95% confidence interval of scores across subjects. Significance of all pairwise differences were computed with Tukey's Honest Significant Difference (HSD) test and $p < 0.01$. Bars with asterisks are significantly greater than chance, computed using a one-sample $t$-test against 0.5 and corrected for multiple comparisons.

**Table 4:** Significant comparisons at the ROI-level that survived post-hoc Tukey correction

| S. No. | ROI 1 | Hemi 1 | Stim 1 | ROI 2 | Hemi 2 | Stim 2 | Mean Difference |
|---|---|---|---|---|---|---|---|
| 1 | sts | rh | IW | psl | rh | IW | 0.221 |
| 2 | sts | rh | W | psl | rh | W | 0.218 |
| 3 | sts | rh | IW | pcg | rh | IW | 0.211 |

| | | | | | | | |
|---|---|---|---|---|---|---|---|
| 4 | pcg | lh | IW | pcg | rh | IW | 0.207 |
| 5 | sts | lh | IW | psl | lh | IW | 0.193 |
| 6 | sts | lh | W | psl | lh | W | 0.189 |
| 7 | sts | rh | W | ifgorb | rh | W | 0.182 |
| 8 | sts | rh | W | ifg | rh | W | 0.179 |
| 9 | pcg | lh | W | pcg | lh | LS | 0.179 |
| 10 | pcg | lh | IW | pcg | lh | LS | 0.179 |
| 11 | sts | lh | LS | psl | lh | LS | 0.175 |
| 12 | sts | lh | LS | pcg | lh | LS | 0.175 |
| 13 | sts | rh | IW | ifgorb | rh | IW | 0.157 |
| 14 | sts | lh | LS | ifgorb | lh | LS | 0.157 |
| 15 | pcg | lh | W | pcg | rh | W | 0.157 |
| 16 | ips | rh | W | vWfa | rh | W | -0.154 |
| 17 | ips | rh | IW | vWfa | rh | IW | -0.154 |
| 18 | sts | rh | W | pcg | rh | W | 0.154 |
| 19 | sts | rh | IW | ifg | rh | IW | 0.15 |
| 20 | sts | rh | IW | sts | rh | LS | 0.146 |
| 21 | ips | rh | LS | vWfa | rh | LS | -0.139 |
| 22 | sts | lh | LS | sts | rh | LS | 0.139 |
| 23 | sts | rh | W | sts | rh | LS | 0.139 |
| 24 | ips | lh | W | vWfa | lh | W | -0.136 |
| 25 | ifg | lh | IW | ifg | rh | IW | 0.136 |
| 26 | psl | lh | W | pcg | lh | W | -0.132 |
| 27 | sts | lh | W | ifgorb | lh | W | 0.129 |
| 28 | sts | lh | IW | ifgorb | lh | IW | 0.129 |

**Table 5:** Cluster × Stimulus comparisons that survived post-hoc Tukey correction

| S. No. | Cluster 1 | Stim 1 | Cluster 2 | Stim 2 | Mean Difference |
|---|---|---|---|---|---|
| 1 | HV | IW | Language | LS | 0.221 |
| 2 | HV | LS | Language | LS | 0.22 |
| 3 | HV | W | Language | LS | 0.207 |
| 4 | EV | LS | Language | LS | 0.179 |
| 5 | EV | IW | Language | LS | 0.173 |
| 6 | HV | IW | Language | W | 0.161 |
| 7 | HV | LS | Language | W | 0.159 |
| 8 | HV | W | Language | W | 0.146 |
| 9 | HV | IW | Language | IW | 0.139 |
| 10 | HV | LS | Language | IW | 0.137 |
| 11 | EV | W | Language | LS | 0.134 |
| 12 | HV | W | Language | IW | 0.125 |
| 13 | EV | LS | Language | W | 0.118 |
| 14 | EV | IW | Language | W | 0.113 |

**Table 6:** Cluster × Hemisphere comparisons that survived post-hoc Tukey correction

| S. No. | Cluster 1 | Hemi 1 | Cluster 2 | Hemi 2 | Mean Difference |
|---|---|---|---|---|---|
| 1 | HV | lh | Language | rh | 0.231 |
| 2 | HV | rh | Language | rh | 0.194 |
| 3 | EV | lh | Language | rh | 0.164 |
| 4 | EV | rh | Language | rh | 0.152 |
| 5 | HV | lh | Language | lh | 0.143 |

### C Functional Connectivity

#### C.1 Agglomerative clustering results for each text condition

In the main paper, the words condition was used to generate the 3 clusters of Early vision, High-level vision, and Language. Here we demonstrate that the same 3 clusters emerge when the analysis is performed on the other text stimuli, with only minor differences at finer-grained levels of clustering below that of the 3 intermediate clusters of interest.

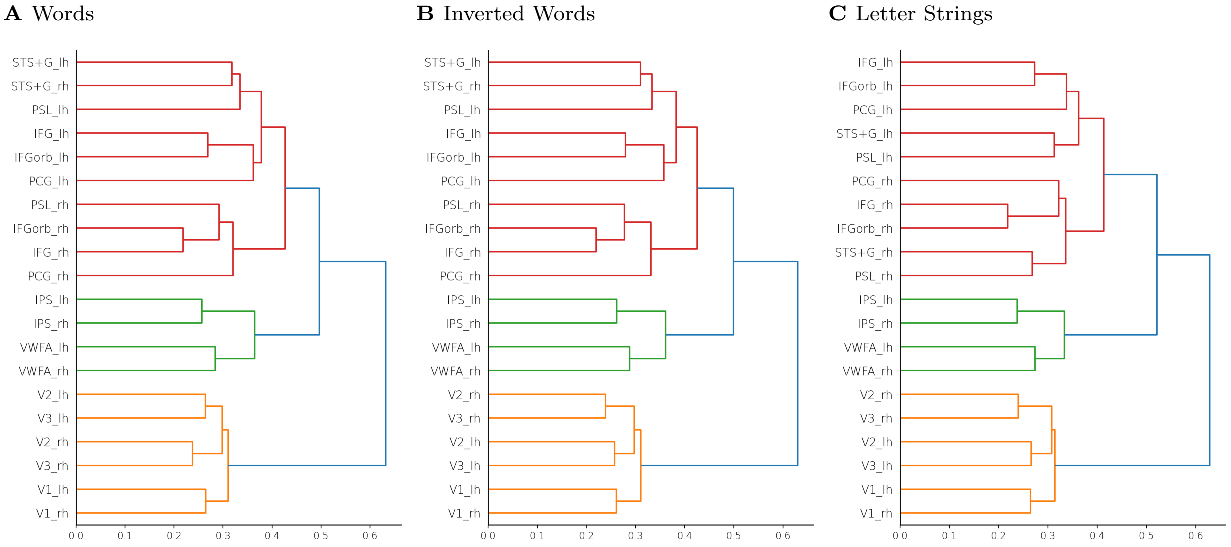


**Figure 9:** Computing dendrograms from mean FC for each of the textual stimulus conditions. The 3 main subnetworks are consistent across each stimulus condition.

#### C.2 Stimulus modulation of FC at the level of ROI-pairs using alternative methods

**A** Left hemisphere

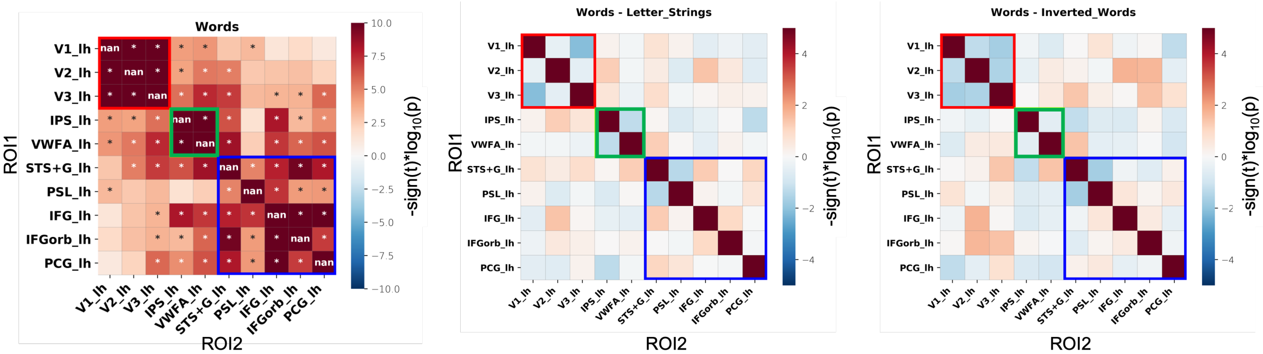


**B** Right hemisphere

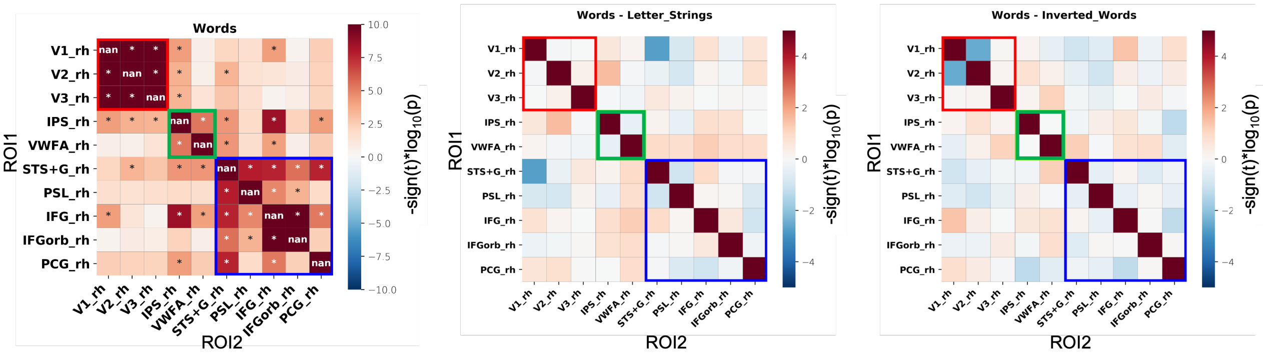


**Figure 10:** Functional correlation (connectivity) of all ROI pairs in the left hemisphere (**A**) and the right hemisphere (**B**) for the three stimulus conditions, using the peak of the HRF signal for the middle time point of each trial. The leftmost panel in **A** and **B** is a one-sample $t$-test of the partial functional correlation between ROI pairs for Words against the null distribution. The middle and rightmost panels in **A** and **B** are two-tailed $t$-tests of the partial functional correlation between ROI pairs for Words compared to Letter Strings and Inverted Words, respectively. Solid red, green and blue boxes surround ROI pairs in the Early vision, High-level vision and Language clusters respectively. Cells with asterisks represent partial functional correlations that survived Bonferroni correction ($p < 0.0005$).

**A** Left hemisphere

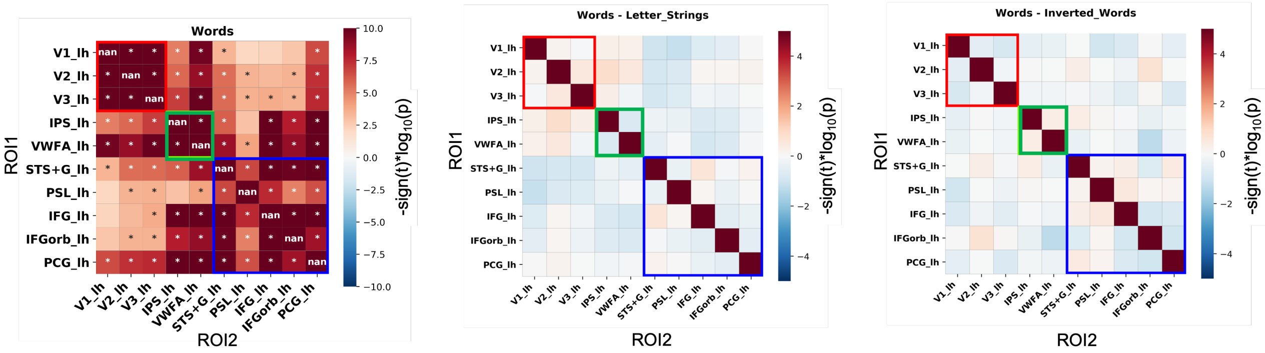


**B** Right hemisphere

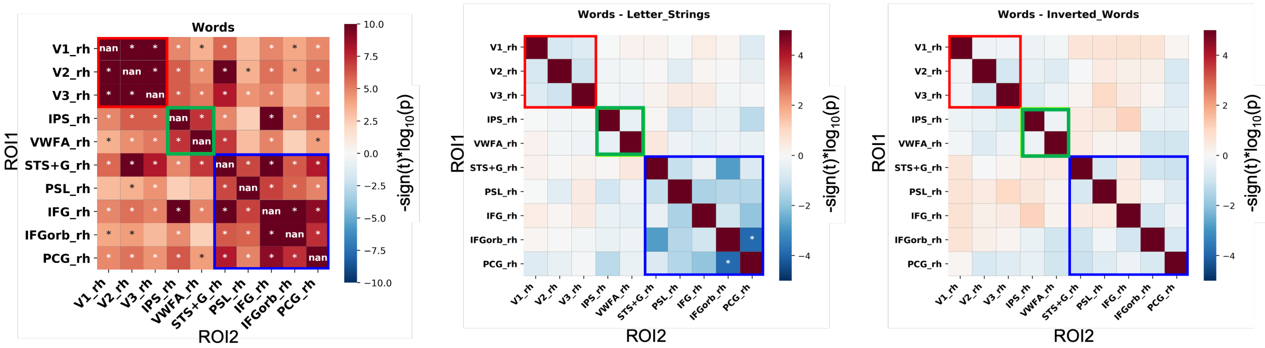


**Figure 11:** Functional correlation (connectivity) of all ROI pairs in the left hemisphere (**A**) and the right hemisphere (**B**) for the three stimulus conditions, using the beta coefficients associated with each stimulus-block obtained from the GLM, across all subjects. The leftmost panel in **A** and **B** is a one-sample $t$-test of the partial functional correlation between ROI pairs for Words against the null distribution. The middle and rightmost panels in **A** and **B** are two-tailed $t$-tests of the partial functional correlation between ROI pairs for Words compared to Letter Strings and Inverted Words, respectively. Solid red, green and blue boxes surround ROI pairs in the Early vision, High-level vision and Language clusters respectively. Cells with asterisks represent partial functional correlations that survived Bonferroni correction ($p < 0.0005$).

#### C.3 Average partial FC of each cluster for the three stimulus types

**Table 7:** Cluster × Stimulus comparisons that survived post-hoc Tukey correction

| S. No. | Cluster 1 | Stim 1 | Cluster 2 | Stim 2 | Mean Difference |
|---|---|---|---|---|---|
| 1 | EV | W | Language | LS | 0.47 |
| 2 | EV | IW | Language | LS | 0.454 |
| 3 | EV | LS | Language | LS | 0.436 |
| 4 | EV | W | Language | W | 0.43 |
| 5 | EV | W | Language | IW | 0.429 |
| 6 | EV | IW | Language | W | 0.413 |
| 7 | EV | IW | Language | IW | 0.412 |
| 8 | EV | LS | Language | W | 0.396 |
| 9 | EV | LS | Language | IW | 0.395 |
| 10 | EV | W | HV | W | 0.355 |
| 11 | EV | IW | HV | W | 0.339 |
| 12 | EV | W | HV | LS | 0.335 |
| 13 | EV | W | HV | IW | 0.323 |
| 14 | EV | LS | HV | W | 0.321 |
| 15 | EV | IW | HV | LS | 0.319 |
| 16 | EV | IW | HV | IW | 0.307 |

| | | | | | |
|---|---|---|---|---|---|
| 17 | EV | LS | HV | LS | 0.301 |
| 18 | EV | LS | HV | IW | 0.289 |
| 19 | HV | IW | Language | LS | 0.147 |
| 20 | HV | LS | Language | LS | 0.135 |
| 21 | HV | W | Language | LS | 0.115 |
| 22 | HV | IW | Language | W | 0.107 |
| 23 | HV | IW | Language | IW | 0.105 |
| 24 | HV | LS | Language | W | 0.095 |
| 25 | HV | LS | Language | IW | 0.094 |
| 26 | HV | W | Language | W | 0.075 |
| 27 | HV | W | Language | IW | 0.074 |

### D Node Strength

#### D.1 Node strength per ROI (using partial functional correlation)

**Table 8:** ROI × Hemisphere comparisons that survived post-hoc Tukey correction

| S. No. | ROI 1 | Hemi 1 | ROI 2 | Hemi 2 | Mean Difference |
|---|---|---|---|---|---|
| 1 | sts | lh | psl | lh | 0.396 |
| 2 | psl | lh | ifg | lh | -0.344 |
| 3 | ips | rh | psl | rh | 0.335 |
| 4 | vwfa | lh | psl | lh | 0.332 |
| 5 | ips | rh | pcg | rh | 0.328 |
| 6 | psl | rh | ifg | rh | -0.319 |
| 7 | ifg | rh | pcg | rh | 0.312 |
| 8 | v1 | lh | psl | lh | 0.308 |
| 9 | vwfa | lh | vwfa | rh | 0.298 |
| 10 | v3 | lh | psl | lh | 0.286 |
| 11 | v2 | rh | psl | rh | 0.276 |
| 12 | ips | lh | psl | lh | 0.274 |
| 13 | sts | rh | psl | rh | 0.274 |
| 14 | v2 | rh | pcg | rh | 0.269 |
| 15 | ips | rh | vwfa | rh | 0.269 |
| 16 | v1 | rh | psl | rh | 0.268 |
| 17 | sts | rh | pcg | rh | 0.267 |
| 18 | v1 | rh | pcg | rh | 0.262 |
| 19 | vwfa | rh | ifg | rh | -0.253 |
| 20 | sts | lh | pcg | lh | 0.25 |
| 21 | psl | lh | ifgorb | lh | -0.247 |
| 22 | v2 | lh | psl | lh | 0.246 |
| 23 | v3 | rh | psl | rh | 0.234 |
| 24 | v3 | rh | pcg | rh | 0.228 |
| 25 | v2 | rh | vwfa | rh | 0.211 |
| 26 | vwfa | rh | sts | rh | -0.209 |
| 27 | v1 | rh | vwfa | rh | 0.203 |
| 28 | ifg | lh | pcg | lh | 0.198 |
| 29 | vwfa | lh | pcg | lh | 0.186 |
| 30 | pcg | lh | pcg | rh | 0.171 |
| 31 | v3 | rh | vwfa | rh | 0.169 |
| 32 | ips | rh | ifgorb | rh | 0.169 |
| 33 | psl | rh | ifgorb | rh | -0.166 |
| 34 | v1 | lh | pcg | lh | 0.162 |
| 35 | ifgorb | rh | pcg | rh | 0.159 |
| 36 | sts | lh | sts | rh | 0.154 |
| 37 | ifg | rh | ifgorb | rh | 0.153 |
| 38 | v2 | lh | sts | lh | -0.15 |
| 39 | sts | lh | ifgorb | lh | 0.149 |
| 40 | psl | lh | pcg | lh | -0.146 |
| 41 | v3 | lh | pcg | lh | 0.14 |
| 42 | ips | lh | pcg | lh | 0.128 |
| 43 | ips | lh | sts | lh | -0.122 |
| 44 | ifgorb | lh | ifgorb | rh | 0.113 |
| 45 | v3 | lh | sts | lh | -0.11 |
| 46 | v2 | rh | ifgorb | rh | 0.11 |
| 47 | sts | rh | ifgorb | rh | 0.108 |
| 48 | v1 | rh | ifgorb | rh | 0.102 |
| 49 | ifgorb | lh | pcg | lh | 0.101 |

| | | | | | |
|---|---|---|---|---|---|
| 50 | v2 | lh | pcg | lh | 0.1 |
| 51 | v3 | rh | ips | rh | -0.1 |
| 52 | vwfa | rh | ifgorb | rh | -0.1 |
| 53 | v2 | lh | ifg | lh | -0.098 |
| 54 | ifg | lh | ifgorb | lh | 0.097 |
| 55 | v1 | lh | sts | lh | -0.088 |
| 56 | v2 | lh | vwfa | lh | -0.086 |
| 57 | vwfa | lh | ifgorb | lh | 0.085 |
| 58 | v3 | lh | v3 | rh | 0.084 |
| 59 | v3 | rh | ifg | rh | -0.084 |
| 60 | v1 | lh | v1 | rh | 0.072 |
| 61 | ips | lh | ifg | lh | -0.07 |
| 62 | v3 | rh | ifgorb | rh | 0.068 |
| 63 | v1 | rh | ips | rh | -0.066 |
| 64 | vwfa | rh | psl | rh | 0.065 |
| 65 | vwfa | lh | sts | lh | -0.064 |
| 66 | v1 | lh | v2 | lh | 0.062 |
| 67 | v1 | lh | ifgorb | lh | 0.061 |
| 68 | ips | rh | sts | rh | 0.061 |
| 69 | vwfa | rh | pcg | rh | 0.059 |
| 70 | ips | lh | vwfa | lh | -0.058 |
| 71 | v2 | rh | ips | rh | -0.058 |
| 72 | ifg | lh | ifg | rh | 0.057 |
| 73 | v3 | lh | ifg | lh | -0.057 |
| 74 | sts | lh | ifg | lh | 0.052 |
| 75 | v1 | rh | ifg | rh | -0.05 |
| 76 | v3 | lh | vwfa | lh | -0.045 |
| 77 | sts | rh | ifg | rh | -0.044 |
| 78 | v2 | rh | v3 | rh | 0.042 |
| 79 | v2 | rh | ifg | rh | -0.042 |
| 80 | v2 | lh | v3 | lh | -0.041 |
| 81 | v3 | lh | ifgorb | lh | 0.04 |
| 82 | v3 | rh | sts | rh | -0.04 |
| 83 | v1 | lh | ifg | lh | -0.036 |
| 84 | v1 | lh | ips | lh | 0.034 |
| 85 | v1 | rh | v3 | rh | 0.034 |
| 86 | psl | lh | psl | rh | 0.032 |
| 87 | ips | lh | ips | rh | -0.029 |
| 88 | v2 | lh | ips | lh | -0.028 |
| 89 | ips | lh | ifgorb | lh | 0.027 |
| 90 | v1 | lh | vwfa | lh | -0.024 |
| 91 | v1 | lh | v3 | lh | 0.021 |
| 92 | ips | rh | ifg | rh | 0.016 |
| 93 | v3 | lh | ips | lh | 0.013 |
| 94 | vwfa | lh | ifg | lh | -0.012 |
| 95 | v1 | rh | v2 | rh | -0.008 |

#### D.2 Node-cluster strength (using partial functional correlation)

**Table 9:** ROI × Cluster × Hemisphere comparisons that survived post-hoc Tukey correction

| S. No. | ROI 1 | Cluster 1 | Hemi 1 | ROI 2 | Cluster 2 | Hemi 2 | Mean Difference |
|---|---|---|---|---|---|---|---|
| 1 | v2 | ev | lh | v2 | lang | lh | 0.245 |
| 2 | v2 | ev | lh | v2 | hv | lh | 0.231 |

| | | | | | | | |
|---|---|---|---|---|---|---|---|
| 3 | v2 | ev | rh | v2 | hv | rh | 0.214 |
| 4 | v2 | ev | rh | v2 | lang | rh | 0.211 |
| 5 | v3 | ev | lh | v3 | lang | lh | 0.202 |
| 6 | v3 | ev | lh | v3 | hv | lh | 0.181 |
| 7 | v1 | ev | lh | v1 | lang | lh | 0.176 |
| 8 | v3 | ev | rh | v3 | lang | rh | 0.167 |
| 9 | v3 | ev | rh | v3 | hv | rh | 0.159 |
| 10 | v1 | ev | lh | v1 | hv | lh | 0.158 |
| 11 | v1 | ev | rh | v1 | lang | rh | 0.135 |
| 12 | v1 | ev | rh | v1 | hv | rh | 0.133 |
| 13 | ifg | ev | lh | ifg | lang | lh | -0.129 |
| 14 | ips | ev | lh | ips | hv | lh | -0.116 |
| 15 | ifgorb | ev | lh | ifgorb | lang | lh | -0.115 |
| 16 | sts | ev | lh | sts | lang | lh | -0.102 |
| 17 | ifg | ev | rh | ifg | lang | rh | -0.09 |
| 18 | vwfa | hv | rh | vwfa | lang | rh | 0.088 |
| 19 | ips | ev | rh | ips | hv | rh | -0.086 |
| 20 | vwfa | ev | rh | vwfa | hv | rh | -0.085 |
| 21 | ifgorb | ev | rh | ifgorb | lang | rh | -0.084 |
| 22 | ifg | ev | lh | ifg | hv | lh | -0.082 |
| 23 | v1 | ev | rh | v2 | ev | rh | -0.081 |
| 24 | ifgorb | hv | rh | ifgorb | lang | rh | -0.08 |
| 25 | ifgorb | hv | lh | ifgorb | lang | lh | -0.073 |
| 26 | psl | ev | lh | psl | lang | lh | -0.072 |
| 27 | vwfa | ev | lh | vwfa | hv | lh | -0.069 |
| 28 | ips | hv | lh | ips | lang | lh | 0.068 |
| 29 | vwfa | hv | lh | vwfa | lang | lh | 0.064 |
| 30 | v1 | ev | lh | v2 | ev | lh | -0.063 |
| 31 | psl | hv | rh | ifg | hv | rh | -0.062 |
| 32 | ifg | ev | rh | ifg | hv | rh | -0.061 |
| 33 | psl | hv | rh | psl | lang | rh | -0.059 |
| 34 | pcg | ev | lh | pcg | lang | lh | -0.058 |
| 35 | psl | ev | rh | psl | lang | rh | -0.057 |
| 36 | ifg | hv | rh | ifgorb | hv | rh | 0.056 |
| 37 | sts | ev | rh | sts | lang | rh | -0.054 |
| 38 | psl | hv | lh | psl | lang | lh | -0.054 |
| 39 | sts | ev | lh | sts | hv | lh | -0.053 |
| 40 | psl | hv | lh | ifg | hv | lh | -0.052 |
| 41 | pcg | ev | rh | pcg | lang | rh | -0.051 |
| 42 | sts | hv | lh | sts | lang | lh | -0.049 |
| 43 | ips | ev | lh | ips | lang | lh | -0.048 |
| 44 | sts | hv | rh | sts | lang | rh | -0.048 |
| 45 | v2 | ev | rh | v3 | ev | rh | 0.048 |
| 46 | ifg | lang | rh | pcg | lang | rh | 0.048 |
| 47 | ifg | hv | lh | ifg | lang | lh | -0.047 |
| 48 | ips | ev | lh | vwfa | ev | lh | -0.047 |
| 49 | ips | hv | rh | ips | lang | rh | 0.046 |
| 50 | psl | lang | lh | ifg | lang | lh | -0.045 |
| 51 | sts | hv | rh | ifg | hv | rh | -0.044 |
| 52 | ifg | lang | lh | pcg | lang | lh | 0.044 |
| 53 | ifgorb | ev | lh | ifgorb | hv | lh | -0.042 |
| 54 | ifgorb | lang | rh | pcg | lang | rh | 0.042 |
| 55 | ips | lang | rh | vwfa | lang | rh | 0.041 |
| 56 | psl | lang | lh | ifgorb | lang | lh | -0.041 |

| | | | | | | | |
|---|---|---|---|---|---|---|---|
| 57 | v1 | ev | lh | v1 | ev | rh | 0.04 |
| 58 | vwfa | lang | lh | vwfa | lang | rh | 0.04 |
| 59 | ips | ev | rh | ips | lang | rh | -0.04 |
| 60 | ifgorb | lang | lh | pcg | lang | lh | 0.04 |
| 61 | sts | hv | lh | psl | hv | lh | 0.037 |
| 62 | pcg | ev | rh | pcg | hv | rh | -0.036 |
| 63 | v3 | ev | lh | v3 | ev | rh | 0.035 |
| 64 | sts | lang | lh | sts | lang | rh | 0.035 |
| 65 | pcg | hv | lh | pcg | lang | lh | -0.035 |
| 66 | vwfa | ev | lh | vwfa | ev | rh | 0.034 |
| 67 | sts | hv | lh | sts | hv | rh | 0.034 |
| 68 | v2 | ev | lh | v3 | ev | lh | 0.034 |
| 69 | v1 | ev | rh | v3 | ev | rh | -0.033 |
| 70 | ifg | hv | rh | pcg | hv | rh | 0.033 |
| 71 | psl | lang | rh | ifg | lang | rh | -0.033 |
| 72 | sts | lang | lh | psl | lang | lh | 0.032 |
| 73 | ifgorb | hv | lh | ifgorb | hv | rh | 0.031 |
| 74 | ifg | hv | lh | pcg | hv | lh | 0.031 |
| 75 | sts | lang | lh | pcg | lang | lh | 0.031 |
| 76 | ifg | hv | rh | ifg | lang | rh | -0.03 |
| 77 | ifg | hv | lh | ifgorb | hv | lh | 0.03 |
| 78 | v1 | ev | lh | v3 | ev | lh | -0.029 |
| 79 | psl | hv | rh | pcg | hv | rh | -0.028 |
| 80 | ifg | ev | lh | pcg | ev | lh | -0.027 |
| 81 | psl | lang | rh | ifgorb | lang | rh | -0.027 |
| 82 | pcg | lang | lh | pcg | lang | rh | 0.026 |
| 83 | sts | lang | rh | ifg | lang | rh | -0.026 |
| 84 | ifgorb | lang | lh | ifgorb | lang | rh | 0.024 |
| 85 | pcg | ev | lh | pcg | hv | lh | -0.023 |
| 86 | ifgorb | hv | rh | pcg | hv | rh | -0.023 |
| 87 | ifg | lang | lh | ifg | lang | rh | 0.022 |
| 88 | psl | hv | lh | ifgorb | hv | lh | -0.022 |
| 89 | sts | lang | rh | pcg | lang | rh | 0.022 |
| 90 | v2 | ev | lh | v2 | ev | rh | 0.021 |
| 91 | v3 | hv | lh | v3 | lang | lh | 0.021 |
| 92 | psl | hv | lh | pcg | hv | lh | -0.02 |
| 93 | sts | lang | rh | ifgorb | lang | rh | -0.02 |
| 94 | pcg | ev | lh | pcg | ev | rh | 0.019 |
| 95 | psl | ev | lh | psl | hv | lh | -0.019 |
| 96 | sts | ev | rh | pcg | ev | rh | 0.019 |
| 97 | v1 | hv | lh | v1 | lang | lh | 0.018 |
| 98 | ips | hv | lh | ips | hv | rh | 0.017 |
| 99 | vwfa | hv | lh | vwfa | hv | rh | 0.017 |
| 100 | ifgorb | ev | lh | pcg | ev | lh | -0.017 |
| 101 | sts | hv | lh | pcg | hv | lh | 0.017 |
| 102 | sts | hv | rh | psl | hv | rh | 0.017 |
| 103 | ifg | ev | lh | ifg | ev | rh | -0.016 |
| 104 | v2 | hv | lh | v3 | hv | lh | -0.016 |
| 105 | v1 | hv | lh | v1 | hv | rh | 0.015 |
| 106 | psl | hv | lh | psl | hv | rh | 0.015 |
| 107 | pcg | hv | rh | pcg | lang | rh | -0.015 |
| 108 | psl | ev | lh | pcg | ev | lh | -0.015 |
| 109 | sts | hv | lh | ifg | hv | lh | -0.015 |
| 110 | sts | hv | lh | ifgorb | hv | lh | 0.015 |

| | | | | | | | |
|---|---|---|---|---|---|---|---|
| 111 | psl | lang | rh | pcg | lang | rh | 0.015 |
| 112 | sts | ev | lh | ifg | ev | lh | 0.014 |
| 113 | v3 | hv | lh | v3 | hv | rh | 0.013 |
| 114 | v2 | lang | lh | v2 | lang | rh | -0.013 |
| 115 | ips | ev | lh | ips | ev | rh | -0.013 |
| 116 | v2 | hv | lh | v2 | lang | lh | 0.013 |
| 117 | sts | ev | lh | pcg | ev | lh | -0.013 |
| 118 | sts | lang | lh | ifg | lang | lh | -0.013 |
| 119 | sts | ev | lh | sts | ev | rh | -0.012 |
| 120 | sts | hv | rh | ifgorb | hv | rh | 0.012 |
| 121 | v1 | hv | lh | v2 | hv | lh | 0.011 |
| 122 | psl | ev | lh | ifg | ev | lh | 0.011 |
| 123 | sts | hv | rh | pcg | hv | rh | -0.011 |
| 124 | psl | lang | lh | psl | lang | rh | 0.01 |
| 125 | sts | ev | rh | ifg | ev | rh | 0.01 |
| 126 | ifg | ev | lh | ifgorb | ev | lh | -0.01 |
| 127 | sts | ev | rh | psl | ev | rh | 0.009 |
| 128 | sts | ev | rh | ifgorb | ev | rh | 0.009 |
| 129 | psl | ev | rh | pcg | ev | rh | 0.009 |
| 130 | ifg | ev | rh | pcg | ev | rh | 0.009 |
| 131 | ifgorb | ev | rh | pcg | ev | rh | 0.009 |
| 132 | sts | lang | lh | ifgorb | lang | lh | -0.009 |
| 133 | ifgorb | ev | lh | ifgorb | ev | rh | -0.008 |
| 134 | v3 | hv | rh | v3 | lang | rh | 0.008 |
| 135 | v2 | hv | rh | v3 | hv | rh | -0.008 |
| 136 | v2 | lang | lh | v3 | lang | lh | -0.008 |
| 137 | v1 | hv | rh | v3 | hv | rh | -0.007 |
| 138 | sts | lang | rh | psl | lang | rh | 0.007 |
| 139 | psl | ev | lh | psl | ev | rh | -0.006 |
| 140 | pcg | hv | lh | pcg | hv | rh | 0.006 |
| 141 | sts | ev | rh | sts | hv | rh | -0.006 |
| 142 | v1 | hv | lh | v3 | hv | lh | -0.006 |
| 143 | v1 | lang | lh | v2 | lang | lh | 0.006 |
| 144 | v1 | lang | rh | v2 | lang | rh | -0.006 |
| 145 | psl | hv | rh | ifgorb | hv | rh | -0.006 |
| 146 | ifg | lang | rh | ifgorb | lang | rh | 0.006 |
| 147 | sts | ev | lh | ifgorb | ev | lh | 0.005 |
| 148 | v2 | hv | lh | v2 | hv | rh | 0.004 |
| 149 | ips | lang | lh | ips | lang | rh | -0.004 |
| 150 | ifg | hv | lh | ifg | hv | rh | 0.004 |
| 151 | v2 | hv | rh | v2 | lang | rh | -0.004 |
| 152 | vwfa | ev | lh | vwfa | lang | lh | -0.004 |
| 153 | ifgorb | ev | rh | ifgorb | hv | rh | -0.004 |
| 154 | v2 | lang | rh | v3 | lang | rh | 0.004 |
| 155 | ifg | lang | lh | ifgorb | lang | lh | 0.004 |
| 156 | ips | lang | lh | vwfa | lang | lh | -0.003 |
| 157 | sts | ev | lh | psl | ev | lh | 0.003 |
| 158 | v1 | hv | rh | v1 | lang | rh | 0.002 |
| 159 | vwfa | ev | rh | vwfa | lang | rh | 0.002 |
| 160 | psl | ev | rh | psl | hv | rh | 0.002 |
| 161 | v1 | lang | lh | v3 | lang | lh | -0.002 |
| 162 | v1 | lang | rh | v3 | lang | rh | -0.002 |
| 163 | psl | ev | lh | ifgorb | ev | lh | 0.002 |
| 164 | ifgorb | hv | lh | pcg | hv | lh | 0.002 |

| | | | | | | | |
|---|---|---|---|---|---|---|---|
| 165 | psl | lang | lh | pcg | lang | lh | -0.001 |
| 166 | v1 | lang | lh | v1 | lang | rh | -9.39e-4 |
| 167 | psl | ev | rh | ifg | ev | rh | 9.29e-4 |
| 168 | ifg | ev | rh | ifgorb | ev | rh | -8.86e-4 |
| 169 | v3 | lang | lh | v3 | lang | rh | -6.27e-4 |
| 170 | ips | ev | rh | vwfa | ev | rh | -5.66e-4 |

#### D.3 Node-cluster strength for Inverted Words and Letter Strings

**A** Node-cluster strength for inverted words

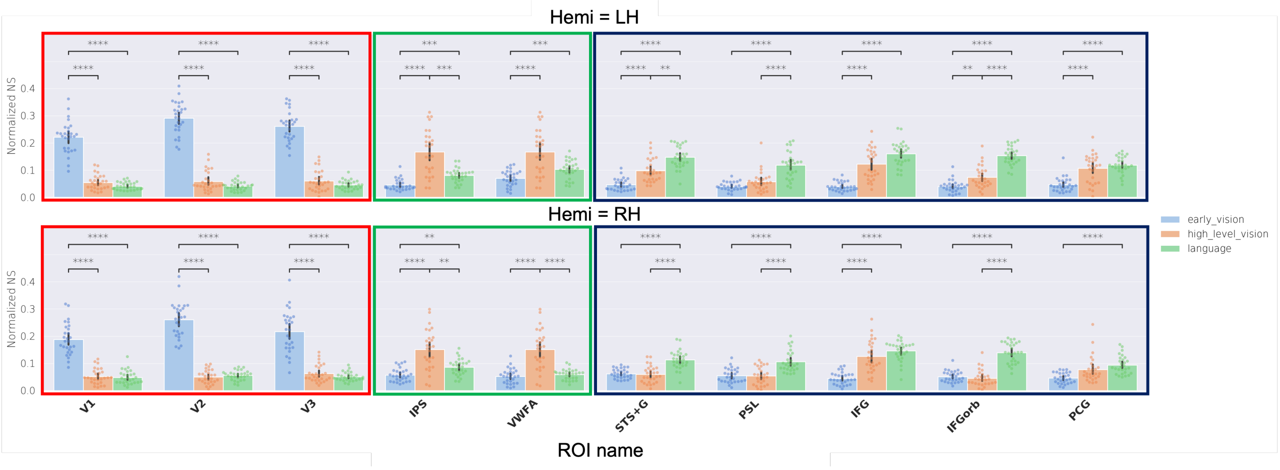


**B** Node-cluster strength for letter strings

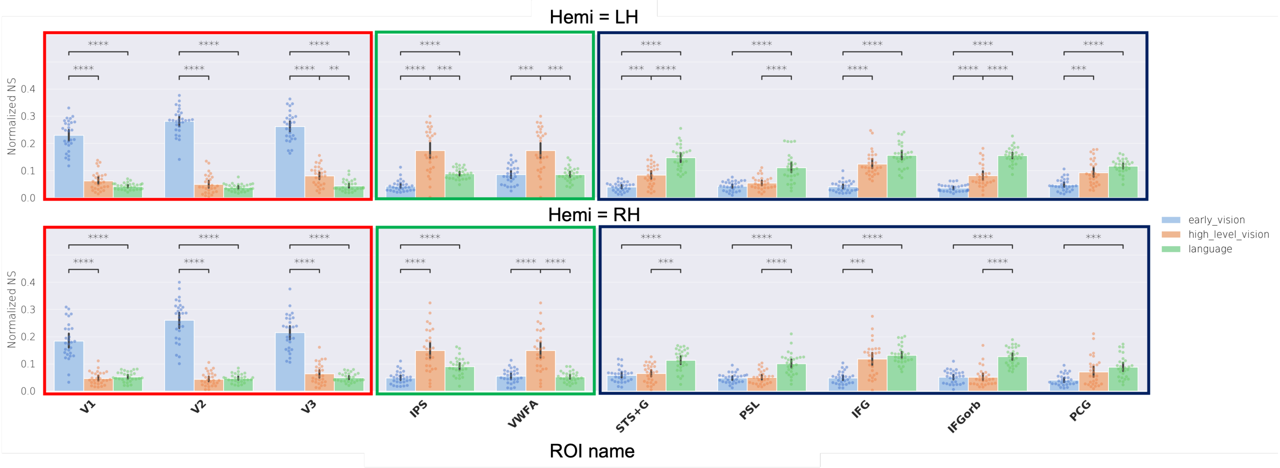


**Figure 12:** Node connectivity to each of the three clusters, in both hemispheres, computed for inverted words and letter strings, and using partial functional correlation. **A.** The normalized connectivity of each ROI with each of the three clusters (Early vision: blue, High-level vision: orange, and Language: green) in the LH (top panel) and the RH (bottom panel) for the Inverted Words stimulus condition. **B.** Normalized connectivity of each ROI to each of the three clusters in the LH (top panel) and the RH (bottom panel) for the Letter Strings stimulus condition. Dots show scores for individual subjects, bars show the average across subjects, and black error lines indicate the 95% confidence interval of the normalized node strength values across subjects. In each plot, horizontal black lines indicate significant comparisons (with $p < 0.05$), and the number of accompanying asterisks indicate the significance (*: $p <= 0.05$, **: $p <= 0.01$, ***: $p <= 0.001$, ****: $p <= 0.0001$).

### E Word list

The words used in the experiment are: [APRIL, BEACH, BIRTH, BLOCK, BLOOD, BREAK, BRIEF, CAUSE, CHAIR, CHECK, CHIEF, CHILD, CLASS, CLEAN, CLOSE, COVER, DAILY, DANCE, DOING, DREAM, DRESS, DRINK, DRIVE, EARTH, EIGHT, EMPTY, ENEMY, EVENT, FIELD, FIGHT, FLOOR, FRONT, GLASS, GROUP, HEART, HORSE, HOTEL, HOUSE, IMAGE, KNIFE, LEAVE, LIGHT, LOCAL, METAL, MIGHT, MONEY, MOUTH, MUSIC, NIGHT, NORTH, OFFER, ORDER, OTHER, PAPER, PARTY, PIECE, PLANT, POINT, PRESS, PRICE, QUIET, RADIO, REACH, RIGHT, RIVER, ROUND, SCORE, SENSE, SERVE, SHAPE, SHARP, SHORE, SIGHT, SLEEP, SMALL, SOUND, SOUTH, SPACE, SPEED, STAGE, STAND, START, STATE, STILL, STORE, STORY, STUDY, SWEET, TABLE, THICK, THING, TODAY, TOTAL, TOUCH, TRADE, TRAIN, TRIAL, TRUTH, UNDER, VALUE, VISIT, VOICE, WATER, WOMAN]. Inverted words were the same, but rotated 180 degrees. Letter strings were random 5-letter consonant strings.
